# Machine learning outcompetes human assessment in identifying eggs of a conspecific brood parasite

**DOI:** 10.1101/2024.11.22.624802

**Authors:** Anna E. Hughes, Lisandrina Mari, Jolyon Troscianko, Václav Jelínek, Tomáš Albrecht, Michal Šulc

## Abstract

Avian brood parasitism provides an exceptional system for studying coevolution. While conspecific brood parasitism (CBP) is more common than interspecific parasitism, it is less studied due to the challenge of detecting parasitic eggs, which closely resemble those of the host. Although molecular genotyping can accurately detect CBP, its high cost has led researchers to explore egg appearance as a more accessible alternative. Barn swallows (*Hirundo rustica*) are considered conspecific brood parasites, but identifying parasitic eggs has traditionally relied on human visual assessment. Here, we used UV-visible photographs of non-parasitized barn swallow clutches and simulated parasitism to compare the accuracy of human assessment with automated methods. In two games, participants and models identified parasitic eggs from six or two options. While humans performed better than chance (72% and 87% accuracy), they still made significant errors. In contrast, the automated supervised model was far more reliable, achieving 95% and 97% accuracy. We think that the model outperformed humans due to its ability to analyse a broader range of visual information, including UV reflectance, which humans cannot perceive. We recommend using supervised models over human assessment for identifying conspecific parasitic eggs and highlight their potential to advance research on evolution of egg colouration.

## Introduction

Avian brood parasitism represents an alternative reproductive strategy wherein parasitic females lay some or all of their eggs in nests belonging to other females. This phenomenon has been documented in 365 species, with 109 species being interspecific obligate brood parasites that consistently lay their eggs in nests of different species (Mann 2017). The interactions between interspecific brood parasites and their hosts have become a major focus for investigating coevolutionary processes in nature (Soler 2017).

Conspecific brood parasitism (CBP) accounts for the remaining 70% of brood parasites and involves individuals occasionally laying eggs in the nests of others within the same species (Yom-Tov and Geffen 2017). Despite its prevalence, CBP remains relatively understudied, largely due to the challenges of detecting it in the wild. Unlike the eggs of obligate brood parasites such as the common cuckoo (*Cuculus canorus*), which, even when highly mimetic, can usually be easily distinguished from host eggs (Mikulica et al. 2017), identifying conspecific parasitic eggs is more difficult because they typically resemble the host’s eggs very closely.

Early studies on CBP relied solely on field observations, such as detecting multiple eggs laid in a single day or noting differences in egg appearance (Yom-Tov and Anderson 1974; YomC:Tov 1980; Møller 1987; Kendra et al. 1988; Brown and Sherman 1989; Jackson 1992). However, later research introduced more precise methods, like protein and genetic fingerprinting, to confirm CBP (e.g. McRae and Burke 1996; Andersson and Åhlund 2001; Griffith et al. 2009). While these genotyping techniques are highly accurate, they remain costly, time-consuming, and technically demanding. As a result, researchers have explored the potential of using egg phenotype and modern analytical methods to identify parasitized clutches and specific parasitic eggs. This non-invasive, low-cost approach also offers the advantage of increasing sample sizes, as genetic fingerprinting may not always be feasible, particularly when predation occurs before blood sampling or when unfertilized eggs without viable DNA are present.

The identification of CBP based on egg phenotype is theoretically possible only in species where individual females lay eggs that are more similar to each other than to those of other females. This pattern, characterized by low variation within clutches and high variation among clutches, appears to be widespread in birds (McRae and Burke 1996; Ornés et al. 2014; Šulc, A.E. Hughes, Troscianko, et al. 2022). Two groups in particular may benefit from this phenomenon: hosts of brood parasites, that can more easily recognize and eject parasitic eggs that differ from their own (Kilner 2006; Cherry and Gosler 2010; Stoddard et al. 2014), and colonially nesting birds, that might otherwise mislay eggs or misdirect incubation and nest defence behaviours (Birkhead et al. 2021; Quach et al. 2021).

The reliability of using egg phenotype to identify parasitized clutches and specific parasitic eggs remains, however, a debated issue. Much of the research on this topic has focused on egg size parameters in waterfowl, which are frequent conspecific brood parasites (Lyon and Eadie 2008). Eadie (1989) proposed an automatic unsupervised method based on maximum Euclidean distance that identifies parasitized clutches by detecting eggs that deviate significantly from the others in the same clutch—more than would typically be expected from a single female’s eggs. His thesis, along with a study by Pöysä et al. (2001), demonstrated that this method is reliable for identifying parasitized clutches of common goldeneyes (*Bucephala clangula*). However, other studies that have replicated this approach in various waterfowl species have reported mixed results, indicating that the method’s reliability vary between species, and they advise using it with caution (Ådahl et al. 2004; Roy et al. 2009; Lemons et al. 2011; Cheng et al. 2016; Petrželková et al. 2017). A more nuanced approach was proposed by Eadie et al. (2010), who suggested categorising clutches into two groups: (1) clutches that can be reliably classified as parasitized or non-parasitized, and (2) clutches where the model’s classification is uncertain. While this more conservative method improved the accuracy, authors still recommend combining its results with observational or molecular techniques for the best outcomes. Recent technological and computational advances have now enabled the application of supervised methods based on machine learning (Christin et al. 2019; Ferreira et al. 2020). Some studies have already used these algorithms, demonstrating that egg phenotypes possess visual identity signals (Gómez et al. 2021), which can be used to identify females that laid them (Šulc, A.E. Hughes, Troscianko, et al. 2022).

Over 30 years ago, Møller’s pioneering studies revealed conspecific brood parasitism (CBP) in the barn swallow (*Hirundo rustica*). He identified this behaviour by noting the appearance of two eggs in active nests (Møller 1987) and by finding eggs in experimental, non-active nests (Møller 1989). Since Møller never directly observed the parasitizing individuals, he relied on egg appearance to identify parasitic eggs and even to speculate on the identity of females that laid them. However, it is well documented that the last eggs laid by various species, including barn swallows, often differ from the other eggs in the clutch (reviewed in Beech et al. 2022), which increases the risk of misidentification when using this method alone (see e.g. McRae 1997; Grønstøl et al. 2006). Therefore, the use of egg appearance for accurately identifying parasitized clutches and specific parasitic eggs in barn swallows still requires further validation.

In this study, we investigated whether egg phenotype could be reliably used to identify parasitized clutches and parasitic eggs in the barn swallow (hereafter swallow), a widely assumed conspecific brood parasite (Møller 1987; 1989; but see Jelínek et al. 2024). Using a labelled dataset of swallow eggs that were molecularly assigned to their genetic mothers, we simulated brood parasitism and tested the ability of human participants, both with and without expert knowledge of swallow egg appearance, to identify foreign (‘parasitic’) eggs within ‘host’ clutches. We also applied analytical techniques to examine phenotypic traits, including size, shape, spotting pattern and colour from ultraviolet-visible photographs and compared the identification accuracy of the unsupervised and supervised tools with that of human participants. The development of these automated methods has the potential to advance research on conspecific brood parasites and the evolution of avian egg colouration.

## Material and methods

### Study population

We collected data during the breeding season in 2020 and 2021 in four farms located in the villages Stará Hlína (49° 02′ 21.4″N, 14° 49′ 06.8″E), Břilice (49° 01′ 14.4″N, 14° 44′ 15.3″E), Lužnice (49°3’25.3"N, 14°46’11.4"E) and Lomnice nad Lužnicí (49°4’7.7"N, 14°42’36.7"E) in southern Bohemia, Czech Republic. In these localities, swallows breed inside cattle barns by building or reusing nests on walls, wooden beams, hanging fluorescent lamps, or in crevices, usually under the ceiling. Swallows start arriving at these breeding sites in late March and females usually start laying eggs in April and May. From May to July we performed four mist-netting sessions and ringed all adults with a unique combination of one aluminium and three plastic colour rings. All individuals were sexed (based on tail length combined with presence/absence of the brood patch and cloacal protuberance), weighed, measured and photographed, and a venipuncture blood sample (∼20 μl) was taken. Most active nests were found before swallows laid the first egg (thanks to the presence of nest lining in the nest) and were subsequently checked every day during the egg-laying period and at 2- to 5-day intervals during nestling stage. At 9 days of age, chicks were ringed and a blood sample (∼10 μl) taken from the jugular vein. All blood samples (from chicks and adults) were stored in 96 % ethanol. We selected 54 clutches, ensuring that a different female laid each clutch and that the same female laid all eggs within a clutch. To verify these assumptions, we genotyped all parents and offspring using 17 microsatellite markers. Alleles were assigned based on raw fragment data processed in Geneious Prime® 2024.0.3 (GraphPad Software LLC d.b.a Geneious). Parentage assignments were conducted with Cervus 3.0.7 (Field Genetics Ltd.). Further details on molecular assignment are available in **Supplementary material** and in Jelínek et al. (2024), which presents a comprehensive maternity analysis for a large dataset spanning 2010 to 2021.

### Egg photography and image analysis

Avian eggs, including those of swallows, reflect visible and also ultraviolet (UV) light (Li et al. 2020). Since birds are known to use UV signals for egg recognition (Honza and Polačiková 2008; Šulc et al. 2016), the UV component may possess important visual identity information. Therefore, we took human-visible and also UV-spectrum photographs of swallow eggs during the first three days of the clutch being complete with a Samsung NX 1000 camera converted to full spectrum, fitted with a Nikkor EL 80 mm lens. Human-visible photographs were taken through a Baader UV-IR blocking filter (Baader Planetarium, Mammendorf, Germany), permitting only visible spectrum light from 420 to 680C:nm and UV photographs were taken with a Baader UV pass filter permitting ultraviolet light from 320 to 380C:nm. All eggs of a given clutch were placed on a dark board and photographed together using a white nylon light tent (50 × 50 × 50 cm, Fomei, China) placed in the shade, at the same angle and from the same distance. Photographing in the shade and inside a diffusive tent limits the variability that could be introduced by changing illumination during the day (see also recommendations by Szala et al. 2023). All photographs were taken in RAW format and referred to two custom-made polytetrafluoroethylene grey standards reflecting 3% and 97% of the UV (300-400nm) and visible (400-700nm) light, respectively. Exposure settings were adjusted accordingly with lighting conditions, yet the ISO value was set constant at 400 and aperture f/8. Image calibration, pattern, colour and shape analysis, and measurements of size were performed in ImageJ software (Schneider et al. 2012) using the Multispectral Image Calibration and Analysis (MICA) Toolbox (Troscianko and Stevens 2015; van den Berg et al. 2020). A scale bar was included in each photo and all images were equally rescaled to the scale of the smallest image (30 pixels/mm).

For colour analysis, we used a custom ImageJ script (Szala 2024) that incorporates functions of MICA toolbox (Troscianko and Stevens 2015). This script is specifically designed to independently measure the brightness and colour of the spots and background of each egg. The script detects spots using the auto local threshold function with the Phansalkar method (Phansalkar et al. 2011) and corrects for uneven illumination with a Gaussian blur (Gómez et al. 2018). We applied a 50-pixel radius for thresholding and a 2048-pixel Gaussian blur. To prevent the dark board on which the eggs were photographed from affecting our measurements, we reduced the selection area around each egg by 3% of its width. The analysis provided averaged reflectance percentage values for the red (R), green (G), blue (B), and ultraviolet (UV) channels, as well as brightness values, for both the spots and the background of each egg. Additionally, using the MICA Toolbox (Troscianko and Stevens 2015), we also calculated pixel proportions across ten luminance levels (ranging from 0 to 1 in 0.1 increments) to capture a more detailed description of the overall luminance of the entire egg.

For pattern analysis, we applied a granularity analysis approach that generates a bandpass energy spectrum across a range of spatial frequencies (Troscianko and Stevens 2015). The pattern energy at each frequency band was quantified as the standard deviation of the filtered image (for further details, see also Šulc et al. 2019). Since pattern energy alone does not differentiate between dark spots on light background and light spots on dark background, we also calculated the skewness of the pattern, which quantifies the asymmetry of the pattern luminance distribution. Additional information and the code for calculating skewness can be found Šulc et al. (2022). Pattern energies and skewness were calculated across the whole egg and also for three egg segments individually (the blunt pole, sharp pole and middle section), to get a measure of how variable the patterning was across the egg.

Additionally, we analysed egg patterning using a custom ImageJ script (Szala 2024) to calculate average spot size, the percentage of the egg covered by spots, and three measures of pattern dispersion. Pattern dispersion parameters offered insights into how the spots are distributed along the egg’s long axis. By analysing pattern coverage for three sections of the egg separately (the blunt pole, sharp pole and middle section), we calculated the mean and the standard deviation (SD) of these coverages, and the coefficient of variation of the pattern dispersion, derived as (SD/mean) × 100.

For shape analysis, we used a tool included in the MICA toolbox (Troscianko and Stevens 2015) and calculated for each egg, the length, maximum width, volume, surface area, ellipse deviation and ellipse aspect ratio (parameter a in Troscianko 2014).

Since swallows usually lay five eggs, we selected photographs of five-egg clutches laid by different females as assessed by molecular analysis (see below). We avoided photographs with low quality photos (blurry or with uneven illumination). The final dataset contained images of 270 swallow eggs laid by 54 females in 54 clutches.

### Preparing variables describing egg appearance

We collected colour, pattern, size and shape data from calibrated photographs for all 270 swallow eggs (54 clutches, each containing 5 eggs). Then, we conducted a principal component analyses (PCA) on different egg features in order to avoid the use of correlated variables in the models. PCA components were selected for the final dataset based on scree plot inspection (Schreiber 2021) and the percentage of variation explained by the selected components are listed below.

#### Colour and luminance data

To perform a PCA for the colour features, we used average R, G, B, UV and brightness values for spots and background for each egg. For luminance, we conducted a PCA using the pixel proportions across ten luminance levels across the entire egg. From these analyses, three colour PCA components (explaining 95% of variance) and one luminance PCA component (explaining 41% of variance) were selected for inclusion in the final dataset.

#### Pattern data

Four PCA components for pattern features were selected for the final dataset. The first two components (explaining 54% of variance) were derived from a PCA that included 12 pattern energy values and 12 skewness values, measured at multiple scales (from 1 to 0.0221 in steps of 1/square root of 2) across the whole egg and within each of three segments of the egg. The remaining two components (explaining 92% of variance) were calculated from a separate PCA on another spot-related variables, including average spot size, percentage of egg surface covered by spots and the mean, standard deviation and coefficient of variance of pattern dispersion.

#### Shape data

The variables entered into the PCA were length, maximum width, volume, surface area, ellipse deviation and aspect ratio. One shape PCA component (explaining 55% of variance) was selected for inclusion into the final dataset.

In total, the final dataset contained 9 PCA components, referred to as egg phenotypic traits, which were used for further analyses.

### Within- and between-clutch variance in egg appearance

For egg appearance to be a reliable indicator of parasitized clutches and parasitic eggs, the variation in egg traits within a single female’s clutch (within-clutch variation) must be lower than the variation between clutches laid by different females (between-clutch variation). To quantify within-clutch variance, we first scaled all phenotypic traits (N = 9) variables across the entire dataset to ensure that no single trait dominates the distance calculations in an algorithm. We then calculated the standard deviation (SD) for each trait within each female’s clutch, and averaged these SD values across all traits, providing an overall variance metric for each female. To quantify between-clutch variance, we calculated the average value of each phenotypic trait for each clutch, effectively creating an ‘average’ egg of each clutch. We then calculated the standard deviation for each phenotypic trait across all clutches and averaged these SDs to generate a metric of between-clutch variance across all traits. To test whether within-clutch variance is indeed lower than between-clutch variance, we performed a one sample t-test, comparing the within-clutch variance metric (N = 54) against the test value representing between-clutch variance.

Additionally, we calculated Beecher’s information statistic (Hs) which quantifies the amount of information an egg’s phenotype conveys about individual identity. This metric is particularly valuable as it allows for comparisons across various studies, species, and signature systems (Beecher 1989; Linhart et al. 2019). This analysis was conducted using the R package ‘*IDmeasurer’* (Linhart et al. 2019). To validate our findings, we compared the results from the real data with a control statistic generated by shuffling the ID labels (Šulc et al. 2022).

### Ranking phenotypic traits by their prediction accuracy

To assess which phenotypic traits are best at predicting parasitic eggs, we fit a random forest model using the R package *‘randomForest’* (Breiman 2001). This model was applied to the entire labelled dataset (comprising all 270 eggs and their associated female identities) to determine the prediction accuracy of each phenotypic trait, measured by mean decrease in accuracy. Mean decrease in accuracy quantifies the reduction in accuracy when a specific variable is excluded, with higher values indicating greater importance for accurate classification. The mean decrease in accuracy values were also used as weights for phenotypic traits in subsequent models aimed at identifying parasitized clutches and individual parasitic eggs. To obtain these weighted phenotypic trait values, each trait was multiplied by its corresponding mean decrease in accuracy value from the random forest model. Using weighted phenotypic traits yielded higher prediction accuracy compared to unweighted traits.

### Identification of parasitized clutches

To assess whether egg appearance can be used to distinguish parasitized clutches from non-parasitized clutches, we adapted Eadie’s (Eadie 1989; Eadie et al. 2010) method based on the maximum Euclidean distance (MED). For each non-parasitized clutch in our dataset (54 clutches in total), we calculated Euclidean distances based on the 9 weighted egg phenotypic traits for each possible egg pair within a clutch (10 pairwise comparisons per clutch of 5 eggs). We then calculated a mean egg difference for each egg by taking the average Euclidean distance of all the egg pairs it was involved in. The value of the most different egg was identified as the MED for that clutch. For parasitized clutches, we created 54 exemplar parasitized clutches by using slides from participant 1 in game 1 (see below for further details), and randomly removed one host egg from each clutch to standardize them to a total of 5 eggs per clutch. We then carried out the same process to calculate the MED for each clutch as for the non-parasitized clutches. We then investigated whether there was a significant difference between the MED for parasitized and non-parasitized clutches using a t-test, as well as plotting the overlap between distributions to assess the utility of this metric for distinguishing between parasitized and non-parasitized clutches.

### Identification of parasitic eggs

We used human assessment and two automatic methods to identify parasitic eggs in barn swallow clutches. All methods of identification used a dataset of 270 eggs laid in 54 clutches and were compared to one another. First, we created calibrated images of all 54 clutches for the purpose of human assessment, using the RGB channels to produce final images of 270 swallow eggs in ImageJ by using the MICA Toolbox (Troscianko and Stevens 2015). These final images were scaled to 10 pixels/mm to match the dimensions of commonly used computer screens. Second, we created parasitized clutches to simulate brood parasitism by adding a swallow egg laid by a different female (the parasitic egg) to a clutch of five eggs laid by the same female the host eggs). This process generated 14,310 unique combinations of parasitized clutches (each of the 54 clutches was parasitized by 265 eggs laid by different females). Then, we randomly selected 1,890 combinations, ensuring that each participant received each of the 54 host clutches with a randomly selected parasitic egg. This approach finally resulted in 1641 unique combinations of parasitized clutches, 114 combinations being duplicated twice and 7 combinations being duplicated three-times.

### Human assessment

To evaluate the human ability to discriminate a parasitic egg from five host eggs, we tested 105 participants, dividing them into three groups based on their experience with bird eggs, specifically those of the barn swallow. The first group, ‘swallow’, consisted of 35 researchers or students with extensive recent experience handling barn swallow eggs. The second group, ‘egg’, included 35 researchers or students with general experience working with wild bird eggs. The final group, ‘bird’, comprised 35 participants with no prior experience with wild bird eggs.

We designed two types of online screen tests (hereafter games) using Psychopy (Peirce 2007) to evaluate participants’ abilities under two different scenarios. Each participant completed both games, with an interval of one to two months between them. Each game consisted of 54 slides displaying parasitized clutches on a grey background.

In the first game, participants were asked to select one egg from a randomly shuffled set of six (complete clutch of five eggs plus one parasitic egg), simulating a scenario in which a researcher encounters a complete clutch without knowledge of the egg-laying order (**Figure 1A**). In the second game, participants were presented with four randomly selected host eggs from one five-egg clutch in the top row, and two eggs below – the remaining fifth host egg and one parasitic egg (**Figure 1B**). Participants were then asked to choose between the two eggs from the bottom row, simulating a scenario where a researcher conducts daily nest checks (a standard ornithological field method) and finds that four eggs have been laid in the normal laying rhythm (i.e., one egg per day), but two eggs were laid on the same day, with one of these two candidate eggs presumed to be parasitic.

**Figure 1.**
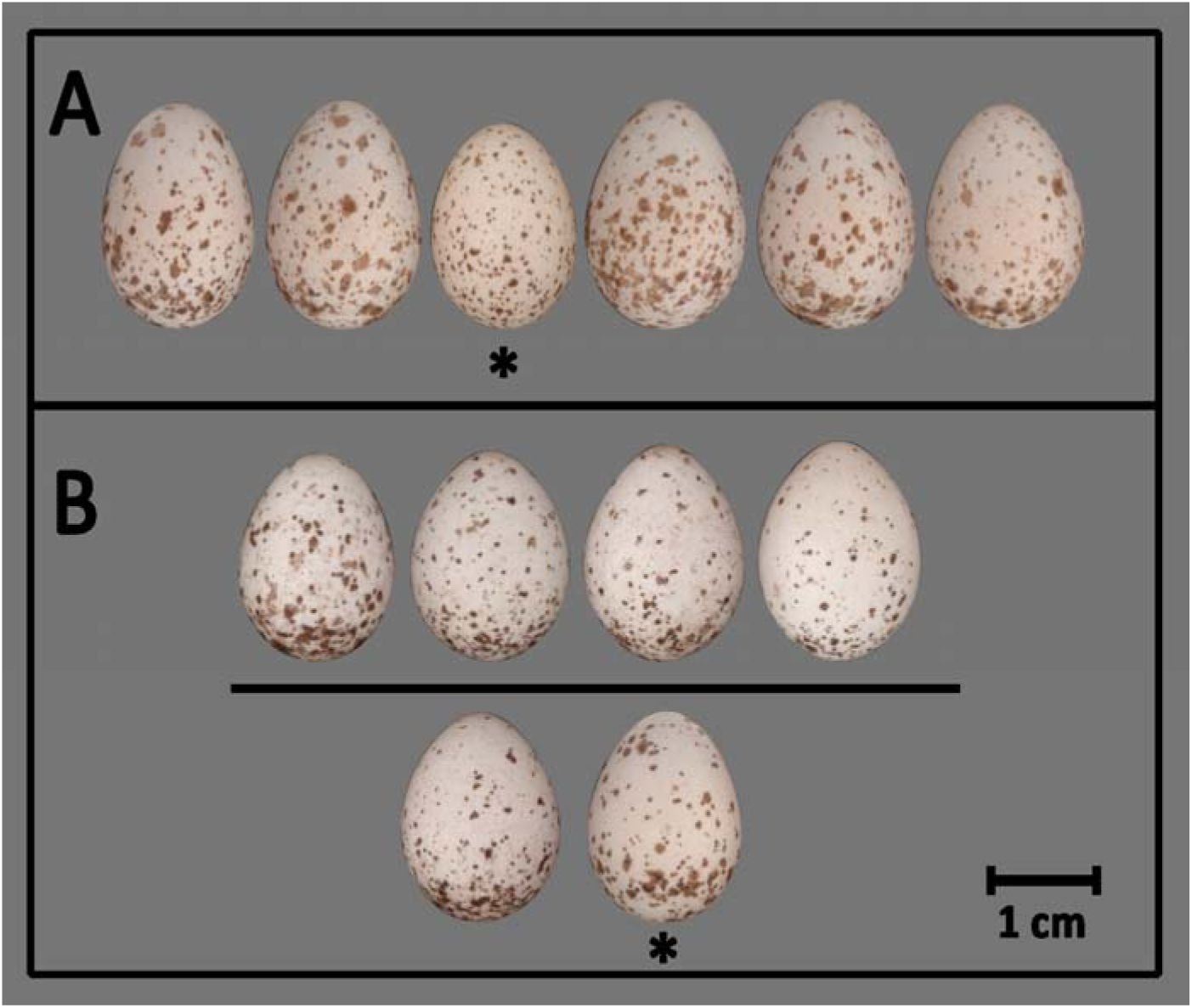
Example slides from the screen games played by participants. Both games consisted of 54 slides, each displaying a six-egg clutch where five eggs were laid by a single ‘host’ female and one egg was laid by a different ‘parasitic’ female. In game 1(A), participants were tasked to visually identify the parasitic egg among the six eggs. In game 2(B), participants were given two egg options in the bottom row and tasked with selecting the parasitic egg, while the four eggs above were known to belong to the host. Asterisks indicate the parasitic eggs, which represent the correct answers.

Every participant within a specific group (swallow, egg, or bird) received a set of different parasitized clutches for the first as well as for the second test. This allowed us to increase our dataset to 1890 combinations of parasitized nests (54 slides for 35 participants = 1890 combinations). Participants with the same number (1-35 in each group) received the same set of parasitized clutches which allowed us to investigate the differences among individual groups. Finally, every participant received the same set of parasitized clutches in their game one and game two so we could test whether and how much participants improved from the first to the second game.

### Automatic assessments

#### Unsupervised classification - Euclidean distance analysis

We carried out an unsupervised classification model for each game. For game 1, this involved calculating a Euclidean distance based on the 9 weighted egg phenotypic traits for each possible egg pair comparison in each parasitized clutch (15 in total, for 6 eggs). We then calculated a mean egg difference for each egg by taking the average of the Euclidean distances of all the egg pairs it was involved in. The parasitic egg assigned by the model was then the egg with the greatest mean egg difference, and we calculated the percentage of times the model was correct across all 1890 clutches presented. For game 2, a similar procedure was followed, except that we only calculated the mean egg differences for the two possible parasitic eggs, and the parasitic egg was selected to be which of these two had the greatest mean egg difference.

### Supervised classification – same/different analysis

We also used an approach where a supervised random forest model was trained to label pairs of eggs as ‘same’ or ‘different’. We used a training dataset consisting of 4000 ‘same’ rows where the two eggs were from the same female, and 4000 ‘different’ rows, where the two eggs were from different females. We then used a ‘leave-one-out’ cross-validation approach (Stone 1974). For each egg in the dataset, the model was trained using a training dataset generated from all other eggs (N=269). In the test phase, we compared the target egg with all other eggs (269 comparisons per egg). Therefore, we obtained 72630 comparisons (270 x 269), two comparisons for each egg pair (one for each egg being a target egg).

For both games, we used the same 1890 combinations of simulated parasitized clutches (always containing one parasitic egg) as used for human assessment. For game 1, we compared every egg pair within a particular parasitized clutch, resulting in 10 comparisons (2x5 comparisons) for each egg. As generating the training dataset involves stochastic processes, we repeated this process 5 times to improve accuracy, yielding a total of 50 comparisons for each egg within the parasitized clutch. The egg with the highest number of ‘different’ model results was considered as the parasitic egg. If the model identified more than one egg as parasitic (i.e. they had the same number of ‘different’ results), we then compared the average Euclidean distances of these eggs from the other eggs of the clutch (see section Unsupervised classification above) and assigned the egg with the greatest distance as parasitic.

For game 2, a similar procedure was followed, except that we compared the two possible parasitic eggs with the remaining four eggs and counted the number of ‘different’ results for the two candidate eggs. Again, the egg with the highest number of ‘different’ results was assigned as parasitic and in case that model resulted in the same numbers, we selected the egg with the higher average Euclidean distance (see section Unsupervised classification above). Finally, for both games we calculated the percentage of times an egg was correctly identified as ‘parasitic’. The full workflow of methods used for the automatic identification of parasitized clutches and parasitic eggs is illustrated in **Figure 2**.

**Figure 2.**
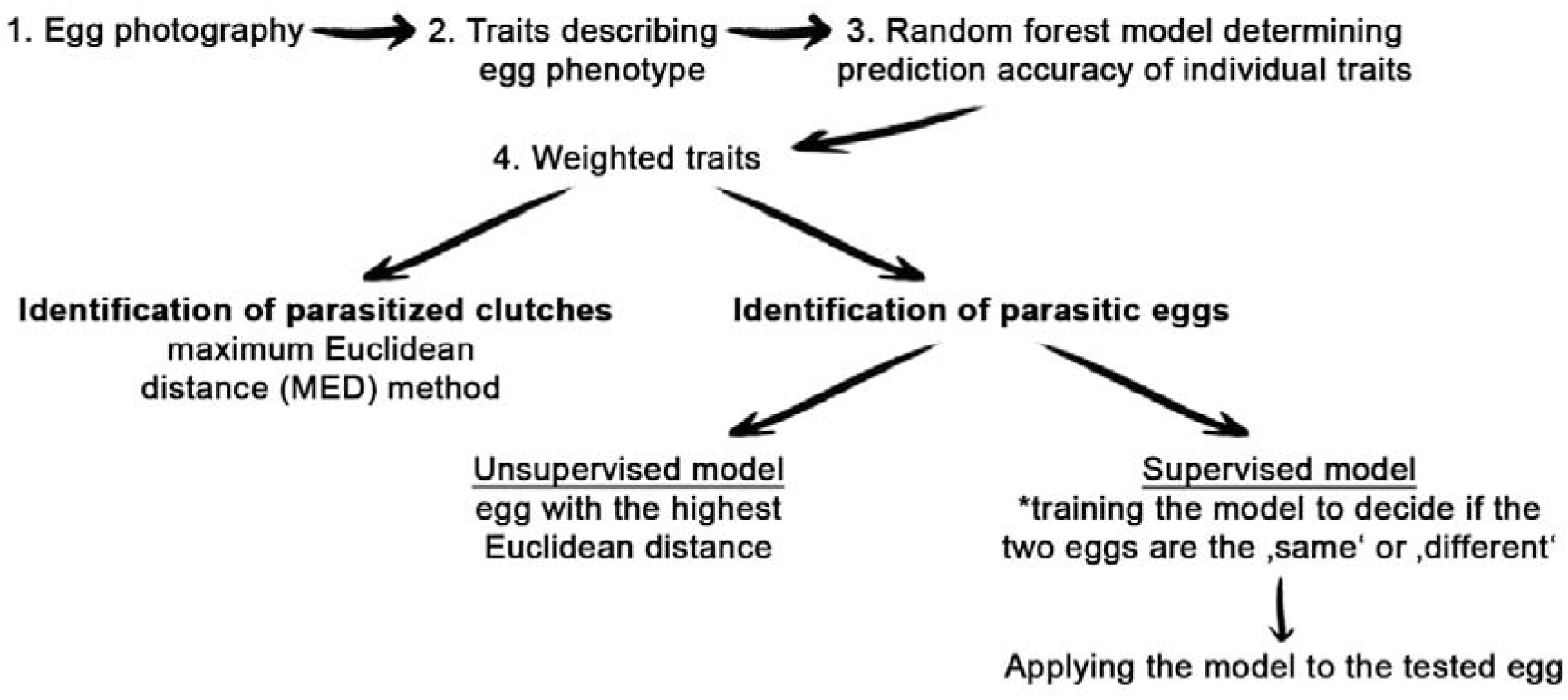
Workflow summary of methods used for the automatic identification of parasitized clutches and parasitic eggs. Asterisk indicate that the model must be trained on the labelled dataset, i.e. with eggs reliably assigned to the individual mothers (e.g., by molecular determination).

## Results

### Within- and between-clutch variance in egg appearance

The mean within-clutch variance was 0.58 (SD = 0.16; range 0.30– 1.06, all clutches can be seen in the **Supplementary material**). Overall, between-clutch variance (mean = 0.82, SD = 0.05) was higher than within-clutch variance (one sample t-test, t = 26.02, df = 53, P < 0.001). Beecher’s information statistic Hs = 0.93 for this dataset, considering only significant variables. (This compares to a control Hs = 0.33, where the ID labels were randomly shuffled). Variation in the egg appearance is also visible in **Figure S1** of the **Supplementary material**.

### Ranking traits for prediction

PC1 for shape was the most important variable for egg classification, and the variables loading onto this PC were the length, maximum width, volume and surface area of the egg. The second most important variable was PC3 for colour, where the spot UV channel, the background UV channel and the spot brightness contrasted with the background visible channels (R and G in particular). The importance of all nine traits used for egg classification is summarised in **Table 1**.

**Table 1.**
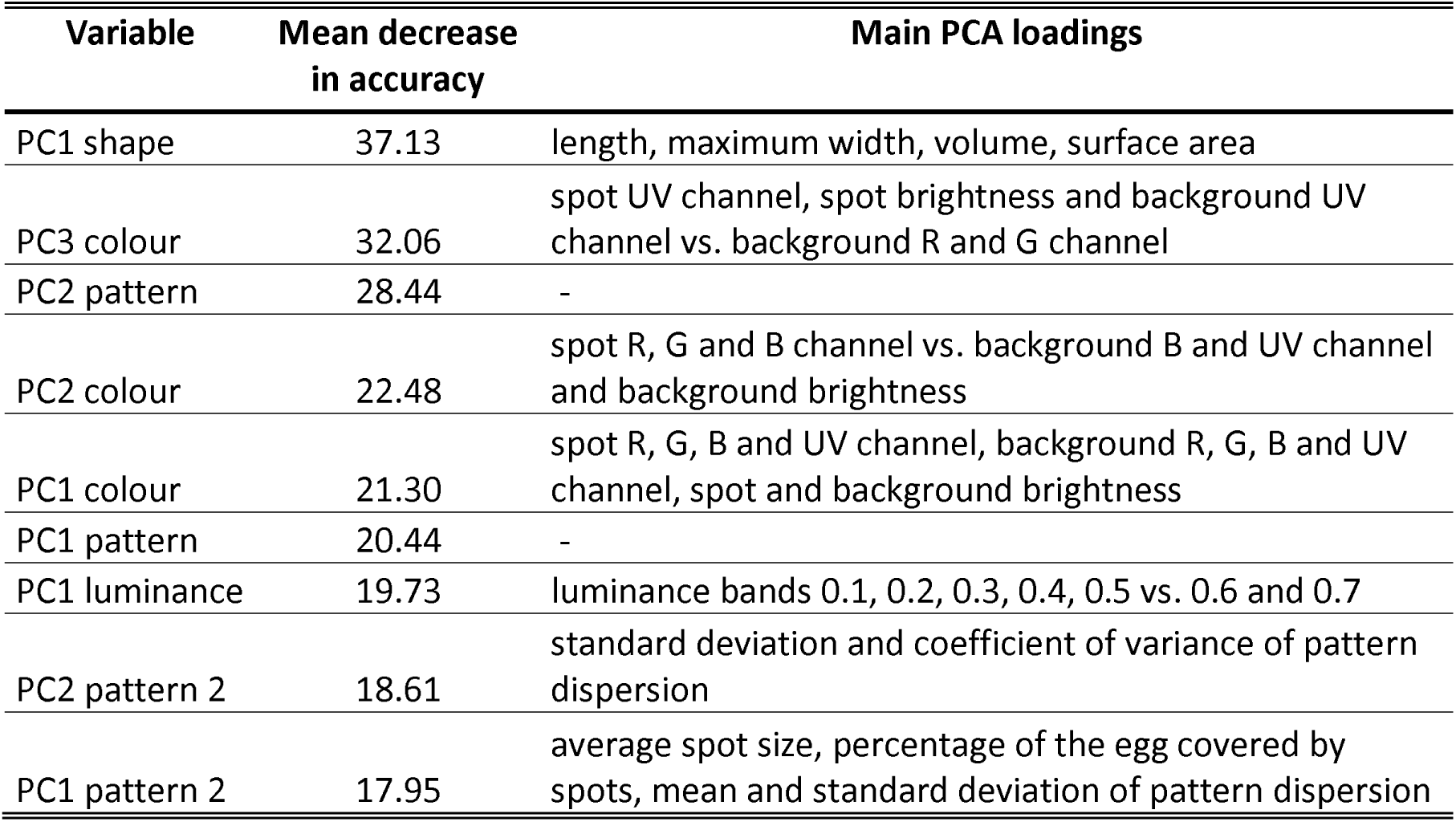
Ranking of trait importance for egg classification by using random forest model. The main PCA loadings are those that were greater than +/– 0.25.

| Variable | Mean decrease in accuracy | Main PCA loadings |
| --- | --- | --- |
| PC1 shape | 37.13 | length, maximum width, volume, surface area |
| PC3 colour | 32.06 | spot UV channel, spot brightness and background UV channel vs. background R and G channel |
| PC2 pattern | 28.44 | - |
| PC2 colour | 22.48 | spot R, G and B channel vs. background B and UV channel and background brightness |
| PC1 colour | 21.30 | spot R, G, B and UV channel, background R, G, B and UV channel, spot and background brightness |
| PC1 pattern | 20.44 | - |
| PC1 luminance | 19.73 | luminance bands 0.1, 0.2, 0.3, 0.4, 0.5 vs. 0.6 and 0.7 |
| PC2 pattern 2 | 18.61 | standard deviation and coefficient of variance of pattern dispersion |
| PC1 pattern 2 | 17.95 | average spot size, percentage of the egg covered by spots, mean and standard deviation of pattern dispersion |

**Table 2.** Summary of identification accuracies in games 1 and 2, achieved by human participants, unsupervised models and supervised models. For the supervised models, reported accuracies include cases where 2% (game 1) and 1% (game 2) of identifications were classified by the unsupervised model.

|  | game 1 | game 2 |
| --- | --- | --- |
| Human assessment | 71.9% | 86.8% |
| Unsupervised model | 62.2% | 86.5% |
| Supervised model | 94.9% | 97.3% |

### Identification of parasitized clutches

The MED was significantly higher in parasitized compared to non-parasitized clutches (t = 4.51, df = 104, p < 0.001; **Figure 3A**), with 96% (52 out of 54) of non-parasitized clutches showing an MED below 350. This suggests that clutches with an MED above this threshold are likely to be parasitized. However, there was a considerable overlap between the MED distributions of parasitized and non-parasitized clutches (**Figure 3B**), and the majority of parasitized clutches (42 out of 54) showed MED values below this threshold. MED on its own therefore cannot reliably distinguish parasitized and non-parasitized clutches.

**Figure 3.**
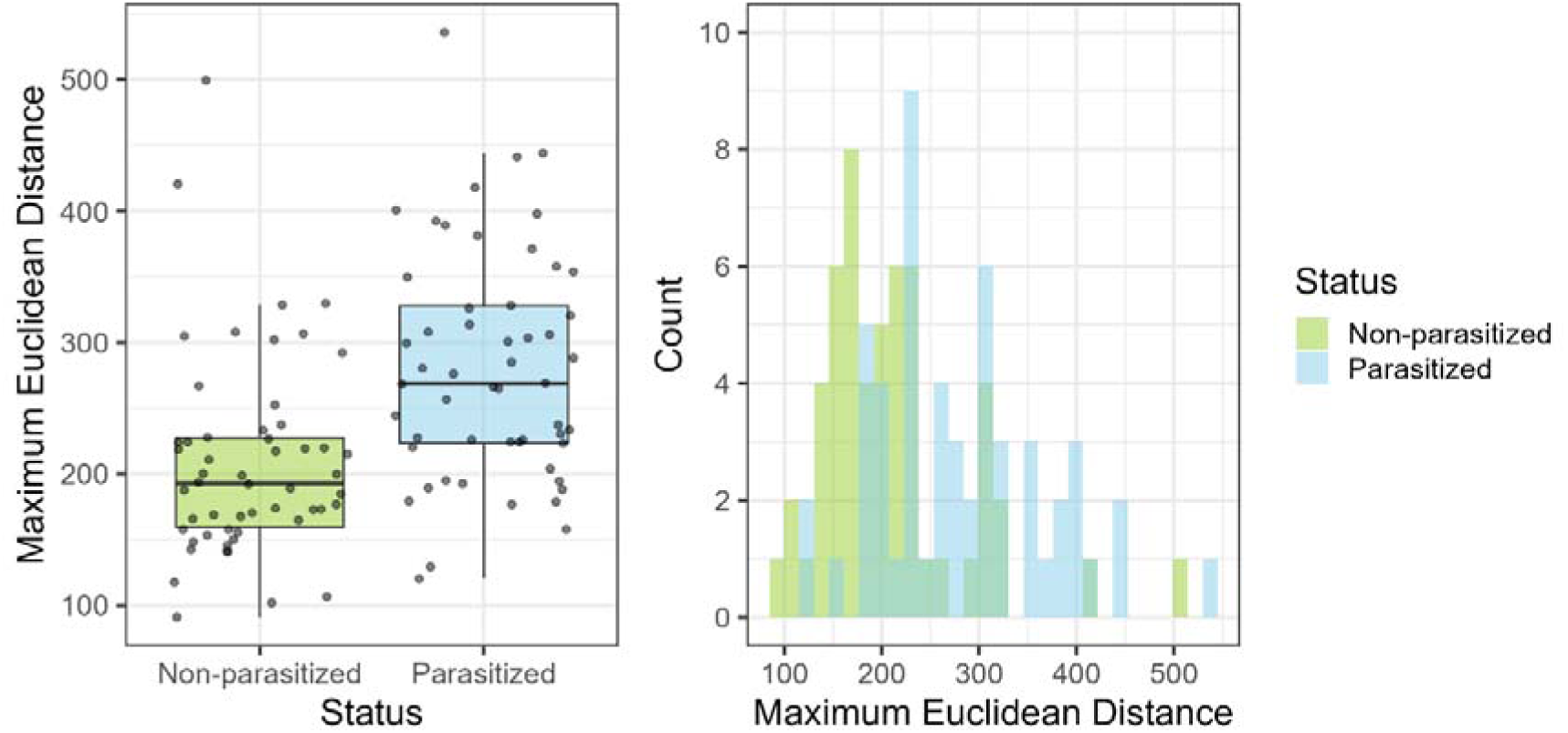
(A) maximum Euclidean distances (MED) calculated for 54 non-parasitized and 54 parasitized clutches. (B) Frequency distribution of MED depicting the substantial overlap between non-parasitized and parasitized clutches.

### Identification of parasitic eggs Human assessment - game 1

Overall, participants performed relatively well on this task, with an average (± SD) of 38.8 ± 6.05 correct egg selections out of 54 trials (ranging from 15 to 49 and corresponding to 71.9% accuracy), significantly above chance (one sample t-test comparing to a mean of 9: t = 50.5, df = 104, p < 0.001; **Figure 4A**). Accuracy did not differ among the three experience groups (one-way ANOVA: F(2,102) = 1.7, p = 0.188), indicating that expertise did not affect performance of the human assessment. The mean time taken on each decision was 10.04s (SD = 13.97).

**Figure 4.**
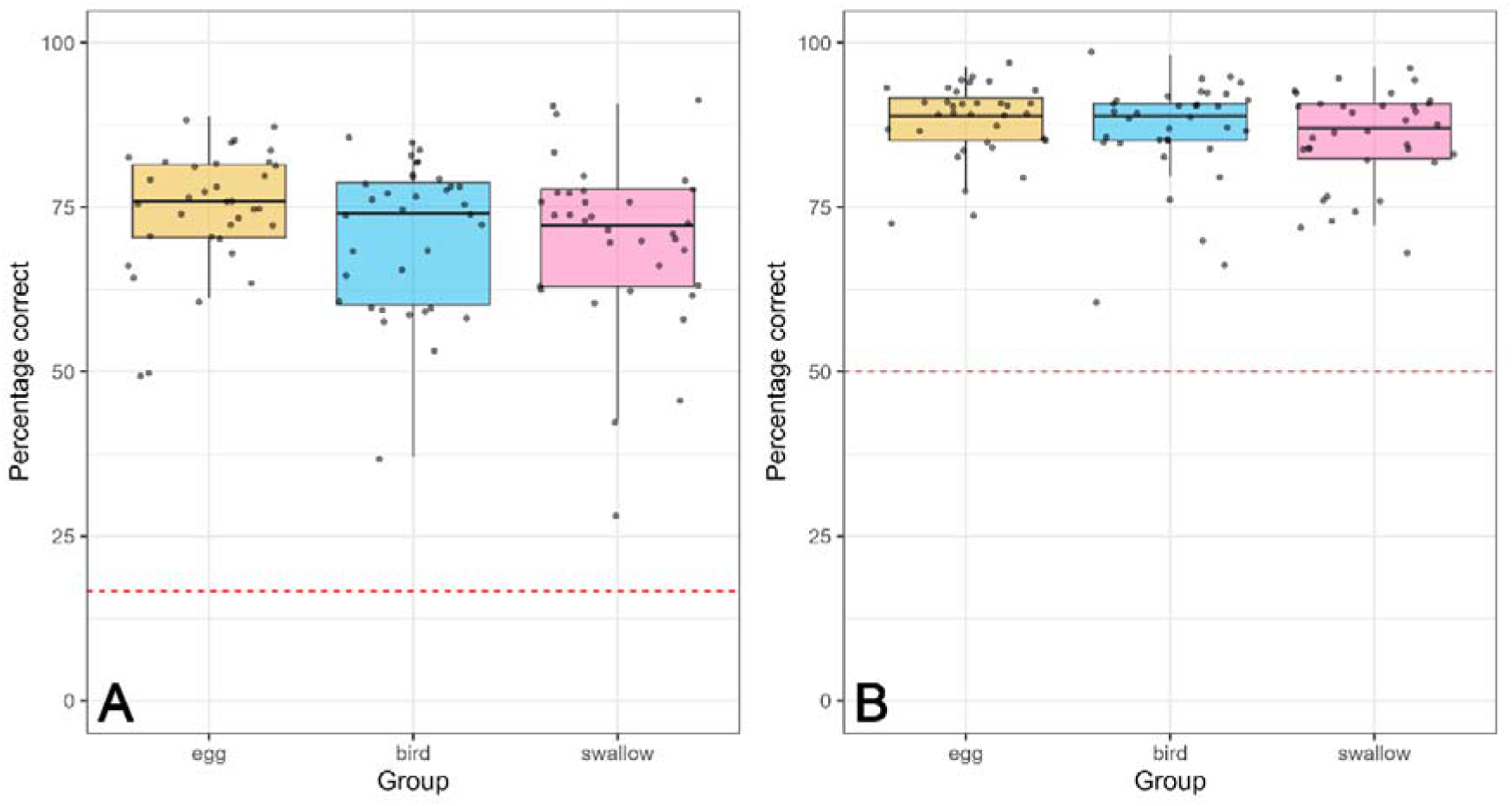
(A) Percentage of correctly assigned parasitic eggs by three differently experienced groups of human participants in the game 1. Since participants were choosing from six-egg options, the red line denotes a level of chance, i.e. 16.7% of correct answers. (B) Percentage of correctly assigned parasitic eggs by three differently experienced groups of human participants in the game 2. Participants were choosing from two-egg options, so the red line denotes a 50% level of chance.

### Human assessment - game 2

Participants performed better on the second game, where they selected between only two eggs, compared to the first game (paired sample t-test: t = -16.851, df = 104, p < 0.001). In the second game, participants achieved a mean accuracy of 46.9 ± 3.8 correct answers out of 54 trials (ranging from 33 to 53 and corresponding to 86.8% accuracy), which was significantly higher than chance (one sample t-test comparing to a mean of 27: t = 53.85, df = 104, p < 0.001; **Figure 4B**). As in the first game, accuracy did not significantly differ among the three experience groups (one way ANOVA: F(2,102) = 1.31, p = 0.274). The mean time taken on each decision was 7.40s (SD = 19.67).

Participants who performed well in game 1 also tended to do well in game 2 (Pearson’s correlation coefficient: r = 0.59, t = 7.45, df = 103, p < 0.001). Additionally, participants who spent more time playing the game generally achieved higher accuracy (average time across games, r = 0.37, t(103) = 4.01, p < 0.001).

### Automatic method - unsupervised classification

In game 1, our unsupervised model based on weighted Euclidean distances predicted the correct parasitic egg in 1175 out of 1890 clutches, corresponding to 62.2% accuracy. This accuracy was significantly lower than accuracy of human participants (t = 5.52, df = 104, p < 0.001). The unsupervised model performed better in game 2 by identifying correctly 1634 parasitic eggs in 1890 clutches, corresponding to an accuracy of 86.5%, which was comparable to human performance (t = 1.13, df = 104, p = 0.26).

### Automatic method - supervised classification

In all possible egg pairwise comparisons, the same-different analysis correctly classified 97% of egg pairs laid by two different females as ‘different’ and only 3% incorrectly as ‘same’ (i.e., incorrectly classified as laid by the same female). However, the model was less effective at identifying egg pairs laid by the same female, which should have been classified as ‘same’, achieving only a 44% success rate and 56% of false alarms.

In game 1, the model correctly identified 1,809 of 1,890 parasitic eggs and in only 71 clutches, another egg was flagged as the parasitic egg. However, due to the high number of false alarms (see above), in 34 out of the 1,809 clutches, one host egg was incorrectly flagged as parasitic alongside the parasitic egg, making it impossible to confidently identify the parasitic egg based on this supervised method. Given the relatively high accuracy of the unsupervised Euclidean distance model in Game 2, we applied this method to these 34 ambiguous cases to determine which of the two candidate parasitic eggs was most distinct from the four host eggs. This approach correctly classified 20 of the 34 parasitic eggs. Overall, this resulted in 1,795 correctly classified parasitic eggs and 85 incorrect classifications, yielding a 94.9% accuracy rate which was significantly higher than human assessment (t = -21.04, df = 104, p < 0.001.

In game 2, the model identified a single egg as parasitic in 1,876 out of 1,890 clutches, of which 1,833 were truly parasitic eggs, and 43 were host eggs. In the remaining 14 clutches, the model did not allow definite decision to be made as both target eggs were flagged as parasitic at the same rate. For these clutches, we used the unsupervised Euclidean distance model, to determine which of the two candidate eggs was most distinct from the four host eggs. This approach correctly classified 6 out of 14 parasitic eggs. Overall, the combination of both methods resulted in 1,839 correct classifications of 1,890, leading to a 97.3% accuracy rate, which significantly outperformed the accuracy of human assessment (t = -15.41, df = 104, p < 0.001).

## Discussion

Our study confirms that eggs laid by the same swallow females are more similar to each other than to eggs laid by different females. This is consistent with previous findings across various species (McRae and Burke 1996; Ornés et al. 2014; Šulc, A. Hughes, et al. 2022). In theory, this pattern should help swallows distinguish foreign eggs from their own, which would be particularly advantageous when defending against brood parasites and/or recognizing own clutches in dense colonies. The former idea has been supported in hosts of interspecific brood parasites, where parasitism exerts strong selective pressure on the evolution of host egg appearance (Stokke et al. 2002; Spottiswoode and Stevens 2011). However, experiments have shown that swallows fail to recognize and reject conspecific or mimetic model eggs when placed among their own (Liang et al. 2013; Šulc, A.E. Hughes, Mari, et al. 2022). This suggests that the level of variation in their eggs is insufficient to trigger a rejection response. Therefore, conspecific brood parasitism is unlikely to be a significant factor driving the reduction of within-clutch variation and/or the increase of between-clutch variation as suggested for hosts of interspecific brood parasites (e.g. Spottiswoode and Stevens 2011; Medina et al. 2016; Caves et al. 2021). Whether this egg variation facilitates recognising their own clutches in dense colonies remains an open question. However, we find this also unlikely, as swallows typically engage in nest guarding behaviour and actively chase away intruders (Møller 1989 and personal observations), including foreign females that could possibly attempt to lay eggs in, or incubate, a foreign clutch. Furthermore, we speculate that the greater variation observed among clutches compared to variation within clutches might be a common phenomenon across bird species, not necessarily driven by selection for egg or clutch recognition. Instead, this pattern likely reflects genetic, developmental, physiological and environmental differences among individual females, which affects the reproductive system, such as shell gland morphology and function. Thus, it is plausible that eggs laid by the same females are more similar in shape and colouration simply due to being produced by the same individual.

Interestingly, the identity signal (Hs) was two times lower for swallow eggs compared to that previously observed in common cuckoo eggs (Šulc, A. Hughes, et al. 2022). This difference was primarily due to between-clutch variance in cuckoos being more than double that of swallows, suggesting that the cuckoo’s strategy of mimicking the egg phenotypes of various host species likely drives this increased variation and identity signal (Stokke et al. 2017; Merondun et al. 2024). This comparison supports the idea that identity signals of bird eggs are strongly influenced by species-specific breeding strategies (Quach et al. 2021).

Our results show little support for using Eadie’s MED method (Eadie 1989; Eadie et al. 2010) to identify parasitized clutches. Although clutches with MED values exceeding 350 had a 96% likelihood of being parasitized, this approach would fail to detect the majority of parasitized clutches (78%) because their MED values fell below this threshold (**Figure 3B**). We suggest that this may be due to the first- or last-laid egg, which often differs noticeably from other eggs in the clutch, as documented in multiple species (e.g. Henriksen 1995; López de Hierro and De Neve 2010, Huo et al. 2018; Mari et al. 2024) including barn swallows (Beech et al. 2022). Our additional analysis of clutches with known laying sequence (N = 32) supports this, showing that both the first- and the last-laid eggs differed the most from the others in the clutch (**Figure S2** in **Supplementary material**). Therefore, we concur with previous studies (McRae 1997; Grønstøl et al. 2006), including the research on closely related American cliff swallows (*Petrochelidon Pyyrhonota*, Brown and Sherman 1989), and advise against relying solely on egg phenotype to distinguish parasitized from non-parasitized clutches. We suggest including additional data, such as observing two eggs laid on the same day, for more accurate identification.

On the other hand, when parasitism is confirmed in a swallow clutch, the combination of low within-clutch and high between-clutch variation enabled us to reach relatively high accuracy in identifying parasitic eggs, especially by the automatic supervised model. In game 1, human participants identified a parasitic egg in a six-egg clutch with 72% accuracy, outperforming the 62% accuracy of an unsupervised Euclidean distance method. As mentioned, the most reliable approach was a supervised "same-different" model, identifying parasitic eggs with the final overall accuracy of 95%. In game 2, where participants selected the parasitic egg between two eggs, mimicking situations with daily nest visits, human accuracy rose to 87%, similarly to the unsupervised method. The supervised model remained the best, successfully identifying 97% of parasitic eggs.

Given the high reliability of the supervised model, we recommend its use for investigating CBP in barn swallows. In contrast, human assessment – while better than chance – made on average errors in 28% and 13% of clutches, respectively, regardless of the participant’s experience. Similar or even worse results were produced for the unsupervised model. Thus, we advise against relying on human assessment or unsupervised models for CBP studies, not only in barn swallows but potentially in other species as well. This caution has already been suggested in previous studies demonstrating insufficient accuracy of human assessment (McRae 1997; Grønstøl et al. 2006; Šulc, A. Hughes, et al. 2022) and unsupervised models (e.g. Cariello et al. 2004; Cheng et al. 2016; Petrželková et al. 2017; Šulc, A. Hughes, et al. 2022). Consequently, we urge caution when interpreting early studies on CBP in barn swallows that relied solely on human assessment to identify parasitic eggs and laying females (Møller 1987; Møller 1989). This is particularly important in the light of recent findings demonstrating that CBP in this species may be less common than previously believed (Jelínek et al. 2024). Overall, our findings show that supervised machine learning models are currently the most effective approach for identifying parasitic eggs.

We believe that the supervised automated method outperformed human participants in identifying parasitic eggs because the model analysed a broader range of visual information in the photographs. In contrast, human perception likely prioritizes certain traits over others, limiting the simultaneous processing of multiple traits to the same extent (Alexander et al. 2019; Hulleman 2020). Additionally, the data used in automated analyses were highly detailed, including information beyond human perception, such as colouration in the UV spectrum. Indeed, UV spot and background colouration were significant contributors to the second most important variable for egg classification. However, we doubt that UV colouration of swallow eggs plays a role in egg or clutch recognition, as our previous findings suggest that UV light barely reaches nests located in buildings (Šulc, A.E. Hughes, Mari, et al. 2022).

The most informative egg traits for the models were related to egg dimensions - length, width, volume and surface area. These traits were also prioritised by human participants during screen games (personal communication with players, and see also Šulc, A. Hughes, et al. 2022). This is encouraging as it suggests that similar analyses could be effective for species with immaculate eggs, where differences in dimensions and background colouration can still be used. Such an approach could improve parasitic egg detection in waterfowl species where previous studies have failed to achieve sufficient accuracy (Ådahl et al. 2004; Roy et al. 2009; Lemons et al. 2011; Cheng et al. 2016; Petrželková et al. 2017). Interestingly, the random forest model indicated that pattern characteristics were less informative, likely because last-laid swallow eggs tend to have lower maculation (Beech et al. 2022), reducing the reliability of this trait for egg identification.

In conclusion, due to high within-clutch variation, egg phenotype alone cannot serve as a reliable cue for identifying parasitized clutches. However, once parasitism is confirmed (e.g., through irregular egg-laying patterns), supervised automated classification has proven highly effective for identifying the parasitic egg. These automated models could be consistently applied to analyse a variety of phenotypic data, enabling objective comparisons across studies and species. Such comparisons would be difficult to achieve with human assessments due to the high level of subjectivity involved within and among individual observers. The only drawback of supervised machine learning models is that they require a labelled dataset with eggs of known identities for training. In the context of egg identification analysis, this requires creating a dataset of clutches with known maternal identities (confirmed in the present study with microsatellite genotyping). By applying these modern analytical methods, future research can explore egg phenotypes across different species to uncover the extent of identity information they convey. This holds great potential to advance our understanding of the evolution of avian egg colouration and its link to interesting behaviours such as brood parasitism and colonial nesting.

## Data availability

All data for these analyses can be found in the Supplementary information.

## Code availability

The code employed in the analyses can be found in the Supplementary information.

## Acknowledgments

We thank our students and colleagues, including K. Bendová, M. Frýbová, M. Janča, K. Míčková, K. Pačandová, L. Pazdera, G. Štětková, O. Tomášek, J. Týblová, J. Záleská, H. Zdobinská and L. Zemanová for their invaluable assistance with fieldwork, as well as all participants who took part in our egg identification games. We are also grateful to K. Szala for providing scripts written that facilitated our analysis of egg coloration and patterning. Finally, our sincere appreciation goes to the managers of the cattle farms that kindly allowed us to conduct our fieldwork on their grounds.

## Author contributions

AEH, MŠ, LM and VJ conceived the ideas and designed methodology; MŠ, VJ and TA collected data; AEH, LM and MŠ performed statistical analyses; AEH, MŠ and LM wrote the initial draft of the manuscript. All authors contributed to the drafts and gave final approval for publication.

## Ethical approval

We declare that all experiments performed for this study were approved by the animal and ethics representatives of The Czech Academy of Sciences and nature conservation authorities (62065/2017-MZE-17214 and MZP/2020/630/964). The fieldwork adhered to the Czech Law on the Protection of Animals against Mistreatment (licence no. CZ03971 and CZ04122).

