## Supplementary material for "Machine learning outcompetes human assessment in identifying eggs of a conspecific brood parasite"

*Detailed methods of molecular assignment of swallow eggs*

We selected 54 clutches, ensuring that a different female laid each clutch, and that the same female laid all eggs within a clutch. To verify these assumptions, we genotyped all parents and offspring using 17 polymorphic microsatellite markers (**Table S1**). DNA was extracted from adult and nestling clotted blood samples and tissue samples (unhatched eggs and dead nestlings) stored in ethanol, using the NucleoSpin Blood kit (Macherey-Nagel, Germany). We followed the manufacturer's instructions, except for the initial proteinase K digestion step which was performed in a homemade digestion solution (20mM EDTA, 50mM Tris pH8, 0,12M NaCl) and was extended to an overnight incubation; and the final elution which was performed in 60ul of buffer AE (10 mM Tris-Cl; 0.5 mM EDTA; pH 9.0). For genotyping, we used a panel of 17 polymorphic microsatellite markers, including two that amplify sex chromosome alleles. We performed two multiplex PCRs per sample (one multiplex mix containing eight and the other one nine fluorescently labelled primer pairs) with the Qiagen Type-It Microsatellite PCR kit (Qiagen). PCRs were conducted in a total reaction volume of 10ul, containing 1ul of DNA (20-100 ug/ul), 1ul of one of the two primer mixes, 5ul of Qiagen Type-It Microsatellite PCR Mastermix and 3ul of PCR-grade water. Conditions were the following: denaturation for five minutes at 95°C, followed by 30 cycles 30s at 94°C, 90s at 50°C for one mix or 52°C for the second, 60s at 72°C, and a final extension step for 45 minutes at 60°C. PCR products were then analysed by capillary electrophoresis. We added 1.5ul of product to 13ul of Hi-Di formamide and GeneScan 500 LIZ dye Size Standard (ThermoFisher Scientific), denatured the mixture for five minutes at 95°C and loaded it on an Applied Biosystems 3130xl BioAnalyzer (ThermoFisher Scientific). Alleles were assigned based on raw fragment data processed in Geneious Prime® 2024.0.3. Bins for each allele were manually created in Geneious, using the shortest and longest PCR fragment lengths of a given amplified allele as the border values of its corresponding bin. All individuals were then genotyped with the binning information. Parentage assignments were conducted using Cervus version 3.0.7 (Field Genetics Ltd.), following the methods of Kalinowski et al. (2007). Parent-pair analysis was carried out using the known sexes of parents.  Further details on the assignments are available in Jelínek et al. (2024), which presents a comprehensive maternity analysis for a large dataset spanning 2010 to 2021.

**Table S1.** *Multiplex compositions, locus details and polymorphism data for the 17 microsatellite loci used in this study.*

Values are based on the genotypes of all individuals. k – number of alleles, C – concentration.

| Loci | Primer sequences (5' - 3') and dye | Mix | C (µM) | Published in | k | marker range (bp) |
|---|---|---|---|---|---|---|
| **TG11-011** | F: 6FAM-ACAAACTAAGTACATCTATATCTGAAG<br>R: TAAATACAGGCAACATTGG | 1 | 1.28 | Dawson et al. 2010 | 14 | 209–233 |
| **HrU5** | F: 6FAM-TCAACAAGTGTCATTAGGTTC<br>R: AACTTAGATAAGGAAGGTATAT | 1 | 0.34 | Primmer et al. 1995 | 41 | 110–144 |
| **2F9** | F: VIC-GCATTTCTGGGCTGTAACAT<br>R: AAAGGACAATGTAATTGGTG | 1 | 0.84 | Heber et al. 2013 | 22 | 78–110 |
| **Hir10** | F: VIC-GGACAAGGGGAGTCTT<br>R: ATTCAGCCAGCCTCTAAT | 1 | 0.18 | Tsyusko et al. 2007 | 15 | 150–202 |
| **3007/3112** | F: PET-TACATACAGGCTCTACTCCT<br>R: CCCCTTCAGGTTCTTTAAAA | 1 | 1 | Ellegren and Fridolfsson 1997 | 4 | 342–354 |
| **Tgu07** | F: PET-CTTCCTGCTATAAGGCACAGG<br>R: AAGTGATCACATTTATTTGAATAT | 1 | 0.4 | Slate et al. 2007 | 17 | 90–124 |
| **TG11-000** | F: PET-TTGCTACCARAATGGAATGT<br>R: TCCTAACCATGAGAAGCAGA | 1 | 1.2 | Dawson et al. 2010 | 17 | 233–265 |
| **Hir11** | F: NED-AACACCTGAAAACCTACAC<br>R: CTTTGAGCAAAATGAGTG | 1 | 0.16 | Tsyusko et al. 2007 | 29 | 164–247 |
| **ADCYAP1_bm** | F: 6FAM-GATGTGAGTAACCAGCCACT<br>R: ATAACACAGGAGCGGTGA | 2 | 0.32 | Steinmeyer et al. 2009 | 25 | 145–184 |
| **Gf16** | F: 6FAM-CCCTTCAGGGCATGAGTGAGG<br>R: ATGTCATGAACTCAACCAACTCC | 2 | 0.18 | Petren 1998 | 14 | 103–130 |
| **IND40** | F: 6FAM-ACCGAAACAACAGAAACAGT<br>R: AGAACGCTAAGTGAATGTCC | 2 | 0.6 | Sefc et al. 2001 | 26 | 222–252 |
| **Hir20** | F: VIC-GAAGTTGGAGAAAGATTAG<br>R: TTATTGCTCTGGGTATGT | 2 | 0.26 | Tsyusko et al. 2007 | 63 | 220–333 |
| **PiJ14-23-CEST** | F: VIC-ATCTGGCATKGAAAACTTGG<br>R: CTCCCTGCACCCCAAAC | 2 | 0.26 | Olano-Marin et al. 2010 | 18 | 156–190 |
| **Z37B** | F: VIC-AACTGGTTGTAGGTATAGTGCAATTATG<br>R: GATTACAAAGCCAATATGGATGC | 2 | 0.28 | Dawson et al. 2015 | 15 | 85–135 |
| **Hir19** | F: PET-GCTCACAACCAGCTAGAC<br>R: ATAGCCACAGGGAAAGTCT | 2 | 0.16 | Tsyusko et al. 2007 | 31 | 130–214 |
| **LEI160** | F: NED-GCAGACAGCCGTTAATATATGCG<br>R: AACCAAAACACAAGCTCTTGCA | 2 | 0.32 | Gibbs et al. 1997 | 16 | 158-188 |
| **Hir22** | F: NED-ATCCGCACCTAATGT<br>R: GATTCACATATCCCATCTAG | 2 | 0.24 | Tsyusko et al. 2007 | 52 | 234–313 |

*Within- and between-clutch variance in egg appearance*

The mean within-clutch variance was 0.58 (SD = 0.16; range 0.30– 1.06, all clutches can be seen at the end of this **Supplementary material**). Overall, between-clutch variance (mean = 0.82, SD = 0.05) was higher than within-clutch variance (one sample t-test, t = 26.02, d.f. = 53, P < 0.001). Beecher's information statistic Hs = 0.93 for this dataset, considering only significant variables. (This compares to a control Hs = 0.33, where the ID labels were randomly shuffled). Variation in the egg appearance is also visible in **Figure S1** where the two most informative variables in the random forest analysis (PC1 for shape and PC3 for colour) are plotted.

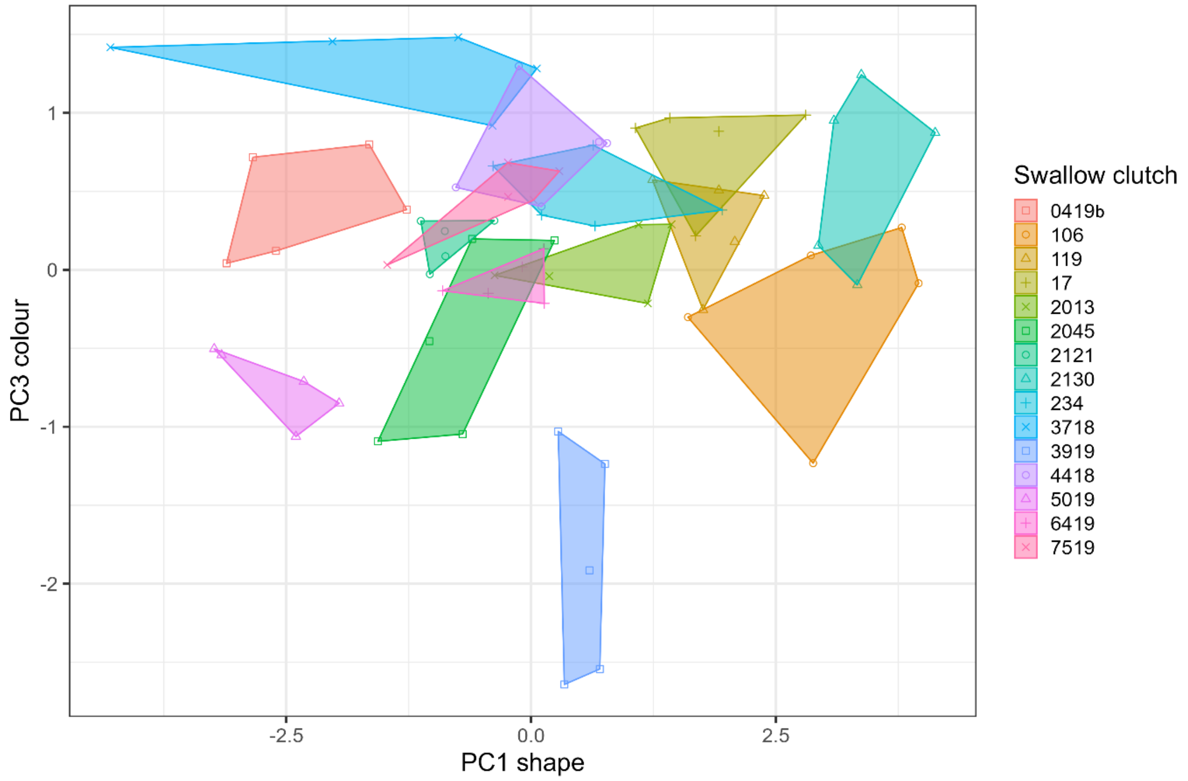


**Figure S1.** *Illustration of within- and between clutch variance in egg traits. Shown are individual egg values for the two most important PC variables (according to the random forest model), grouped by swallow clutch ID with 15 clutches randomly selected. PC1 shape indicates egg dimensions as length, width and volume and PC3 colour indicates the contrast between spot UV channel, background UV channel and the spot brightness with the background visible channels (R and G in particular). For further details, see* ***Table 1*** *in the main manuscript.*

*Effect of laying order on within-clutch egg dissimilarity*

We had complete laying sequence data for 32 of 54 clutches (i.e. from the first to the fifth egg). For each egg in these clutches, we calculated the average Euclidean distance to the other eggs within the same clutch, based on nine weighted traits (see main manuscript). To examine the effect of laying order on egg dissimilarity within clutches, we compared the average Euclidean distances across laying order groups using repeated measures ANOVA and pairwise t-tests. We hypothesized that the last-laid egg would show the highest dissimilarity, as previous studies have documented this pattern across various species (Henriksen 1995; López de Hierro and De Neve 2010; Huo et al. 2018; Mari et al. 2024), including barn swallows (Beech et al. 2022).

Our results confirmed that laying order significantly affects egg phenotype (repeated measures ANOVA, F(2.89, 89.67) = 5.54, p = 0.002, **Figure S2**). Specifically, last-laid eggs differed significantly from the second, third and fourth eggs (p < 0.01) and first-laid eggs also showed significant differences from the second and third eggs (p < 0.05).

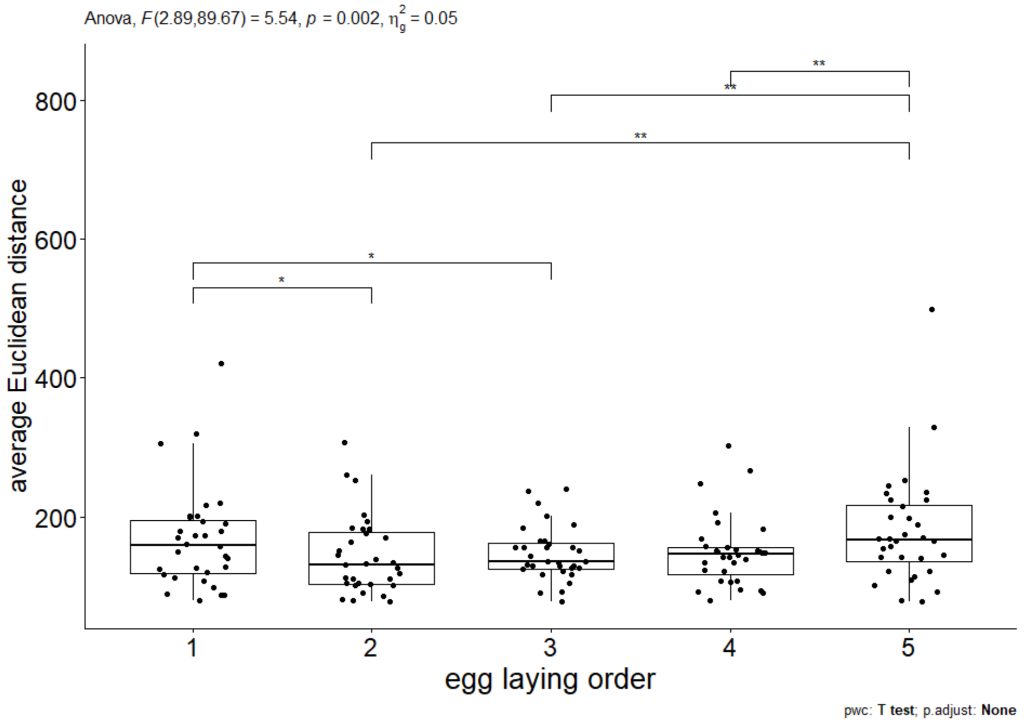


**Figure S2.** *Pairwise comparison of the average Euclidean distance of individual barn swallow eggs to other eggs within the same clutch. Lines indicate significant pairwise differences, with * representing p < 0.05 and ** indicating p < 0.01.*

### *Calibrated images of barn swallow eggs*

The images display 270 barn swallow eggs laid by 54 females in 54 clutches, with each row representing a different clutch. Stars next to the clutch numbers indicate clutches with known laying sequence, arranged from the first to the fifth egg (left to right). The images were calibrated and generated using RGB channels in ImageJ, by using the MICA Toolbox (Troscianko and Stevens 2015). For additional details in this process, please refer to the Material and Methods section in the main article.

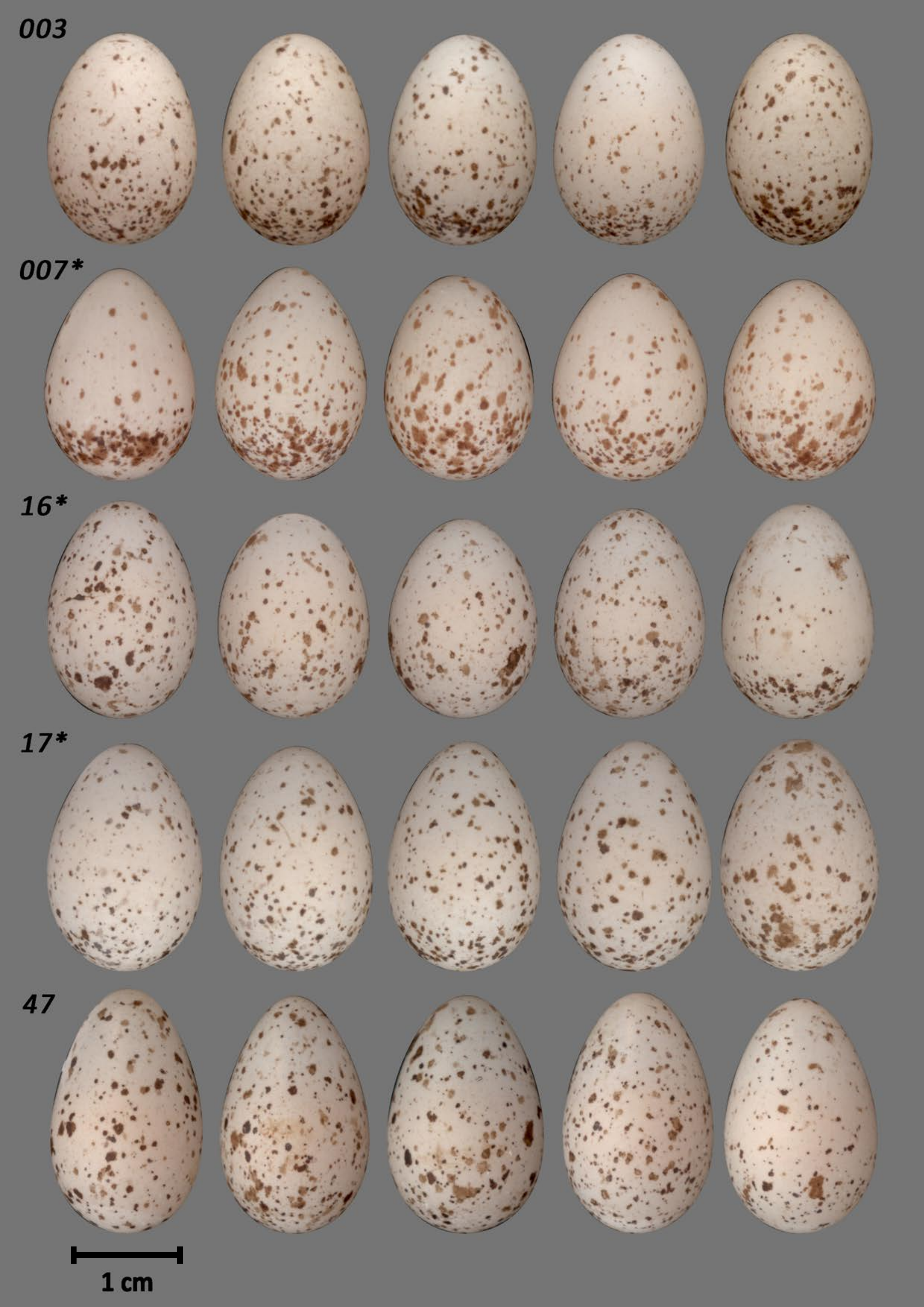
003
007*
16*
17*
47
1 cm

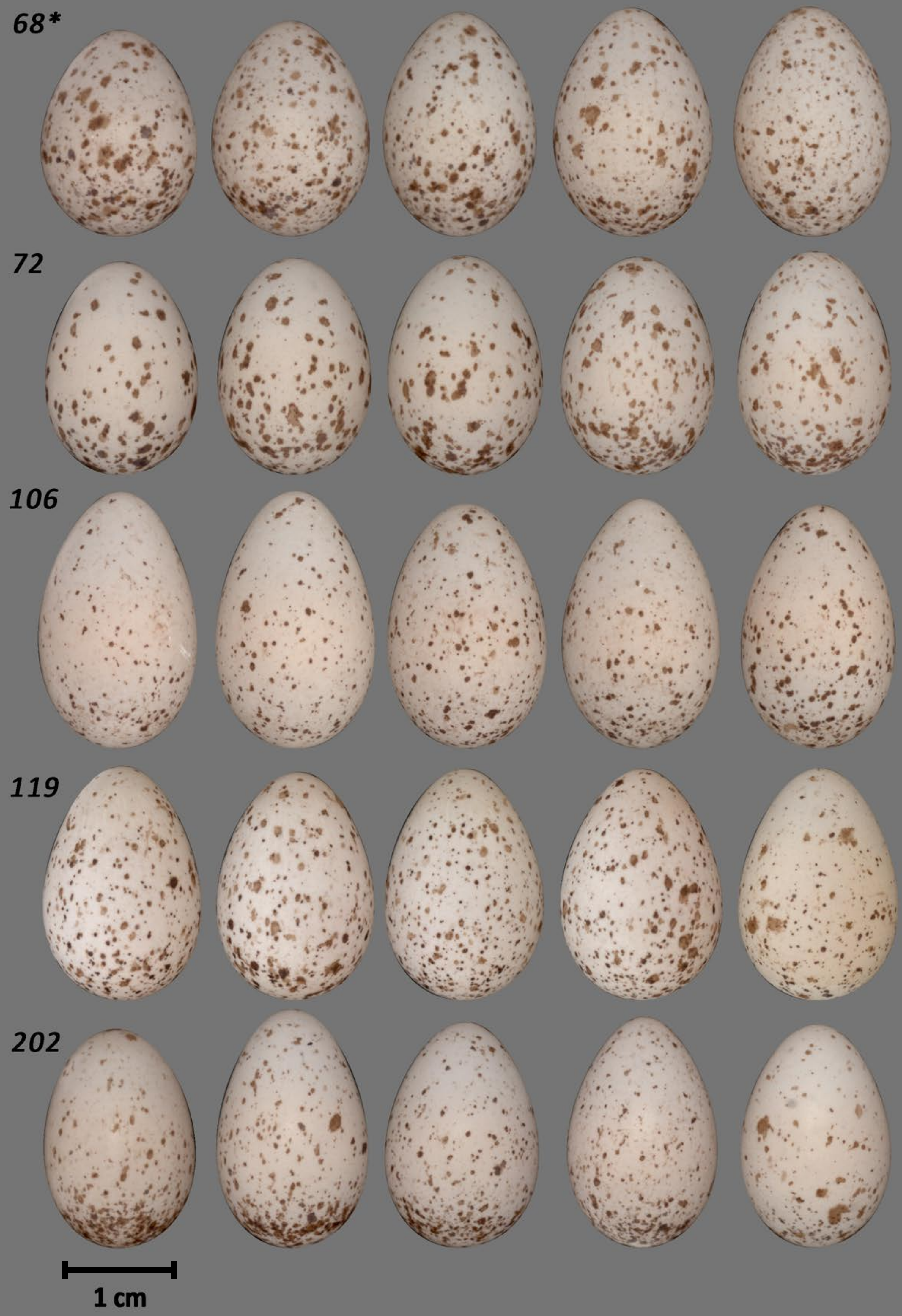
68*
72
106
119
202
1 cm

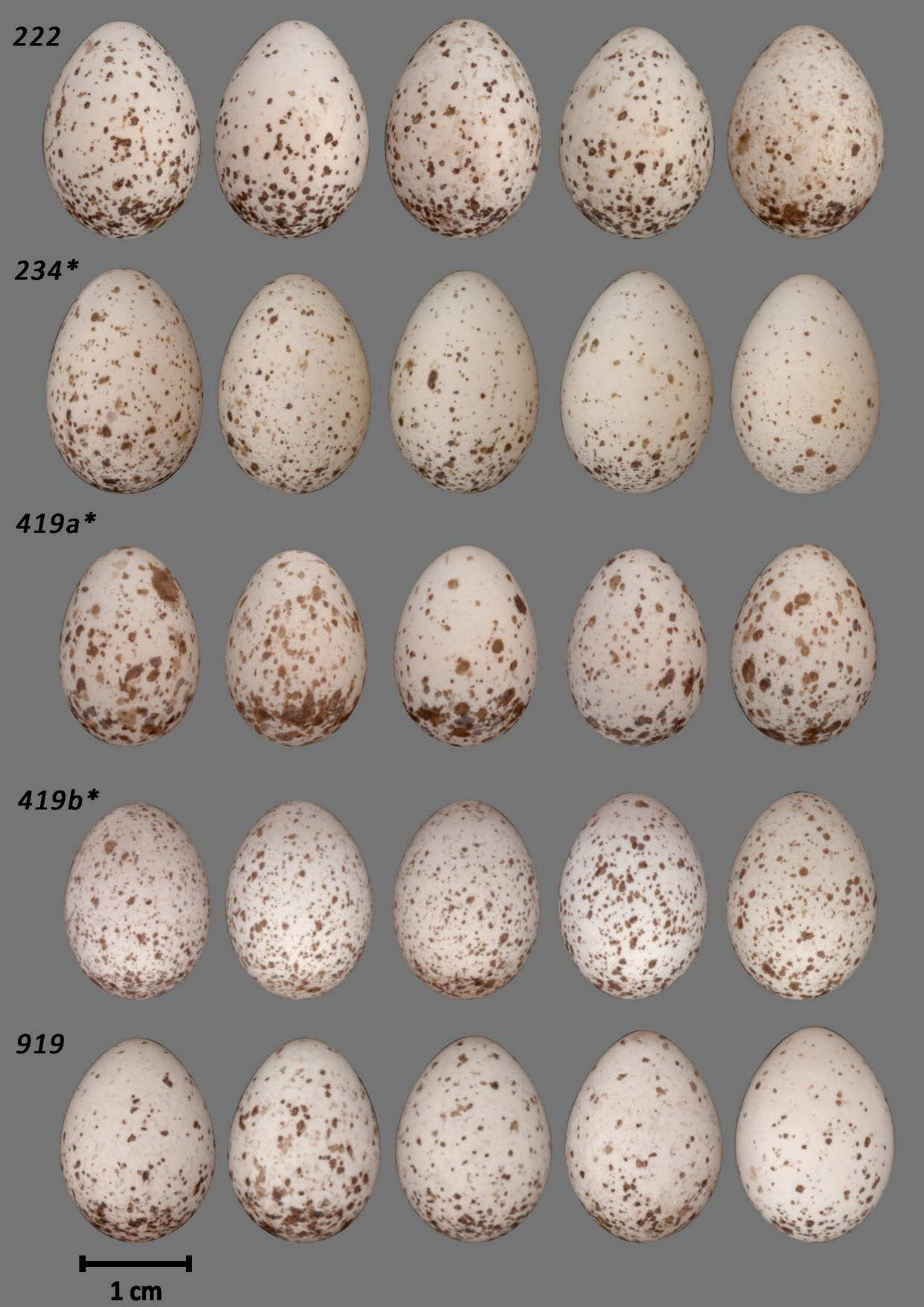
222
234*
419a*
419b*
919
1 cm

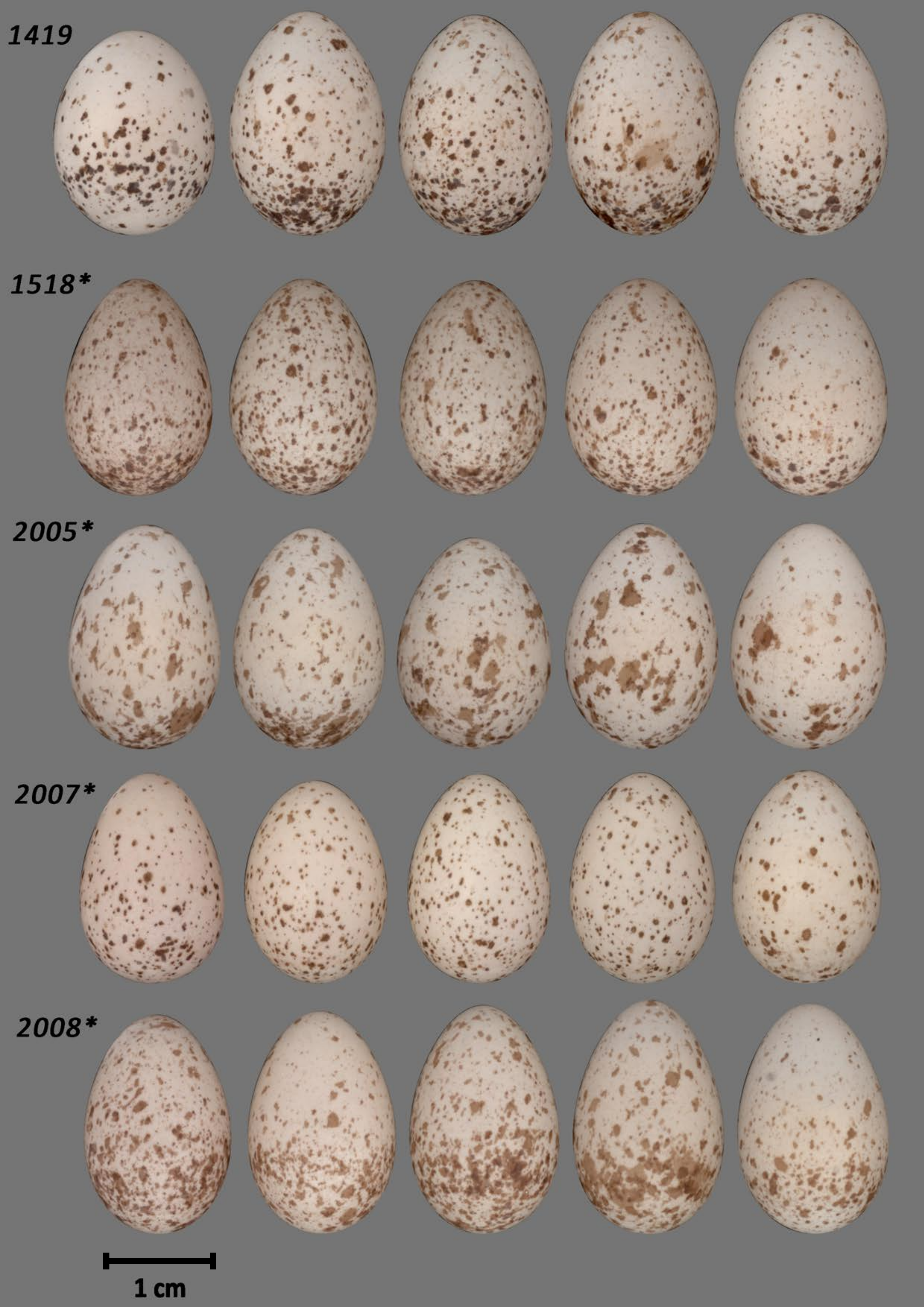
1419
1518*
2005*
2007*
2008*
1 cm

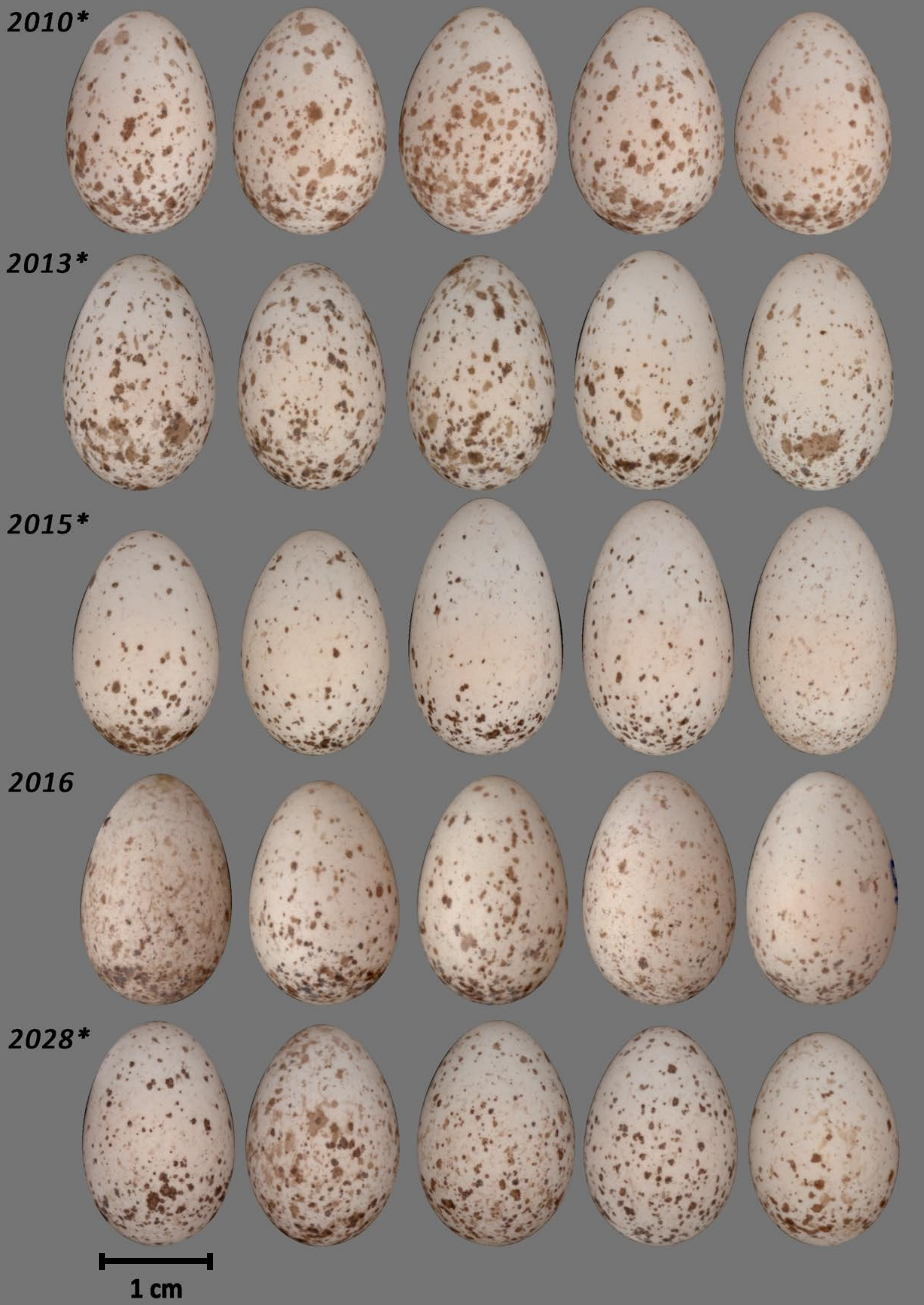
2010*
2013*
2015*
2016
2028*
1 cm

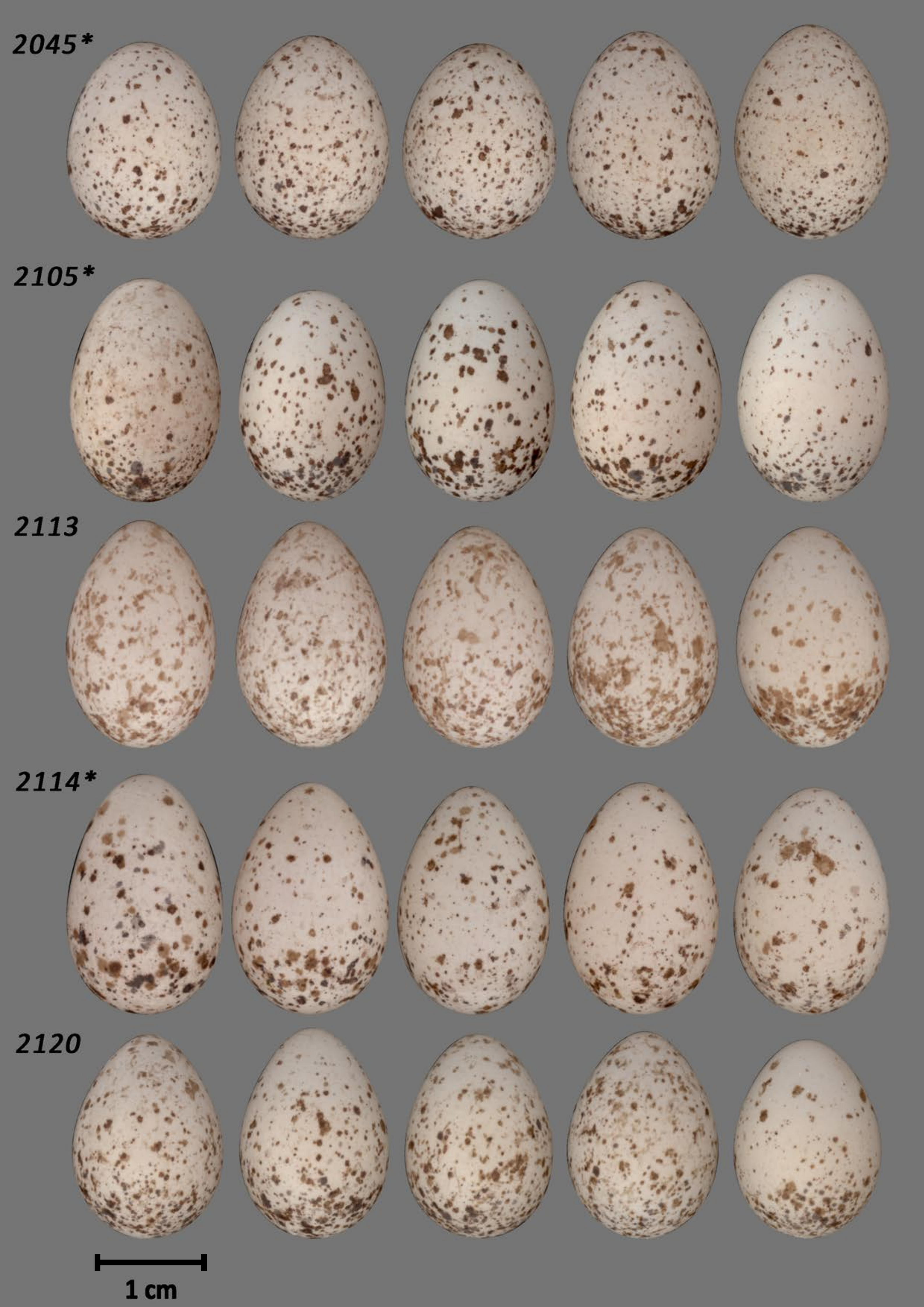
2045*
2105*
2113
2114*
2120
1 cm

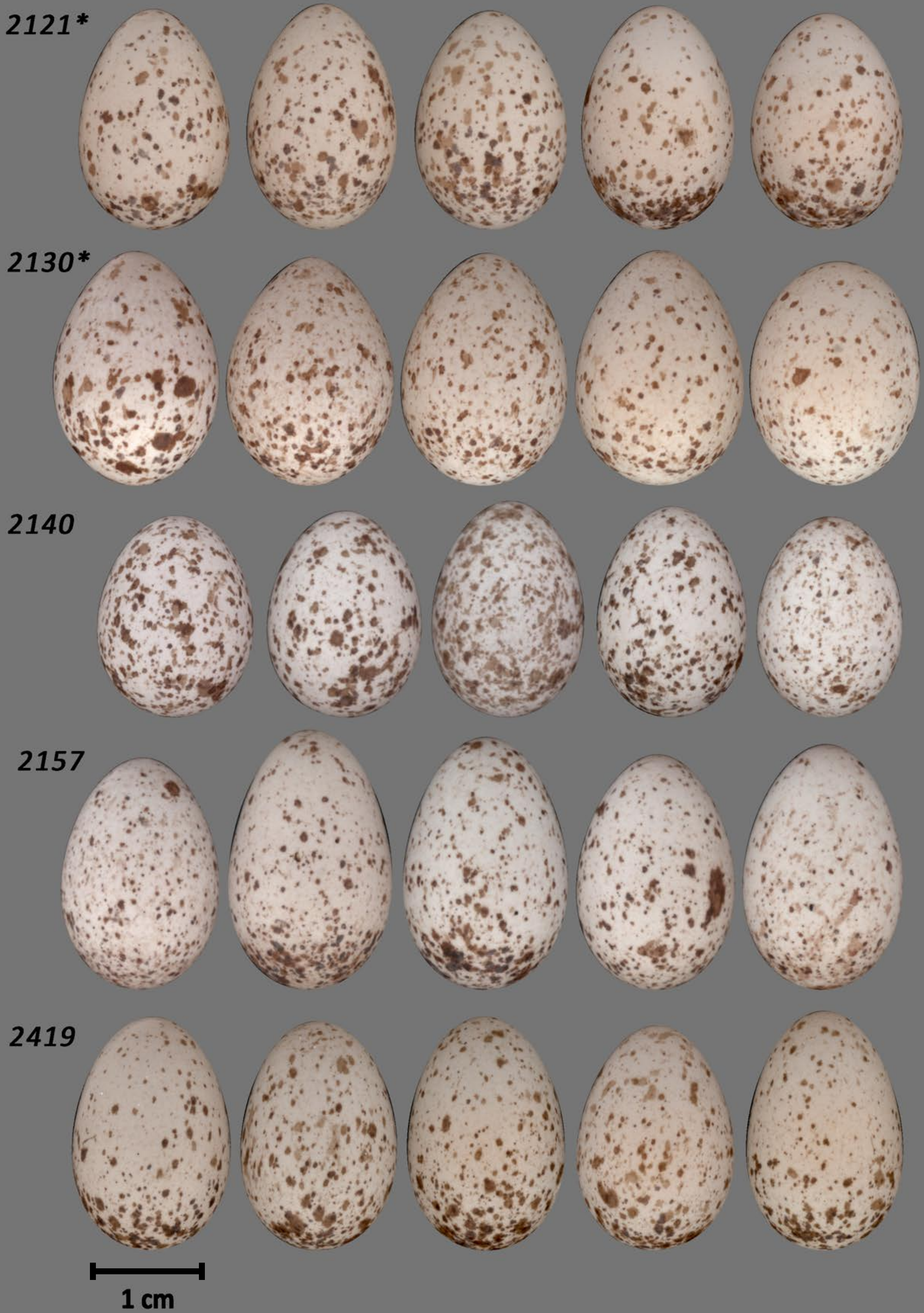
2121*
2130*
2140
2157
2419
1 cm

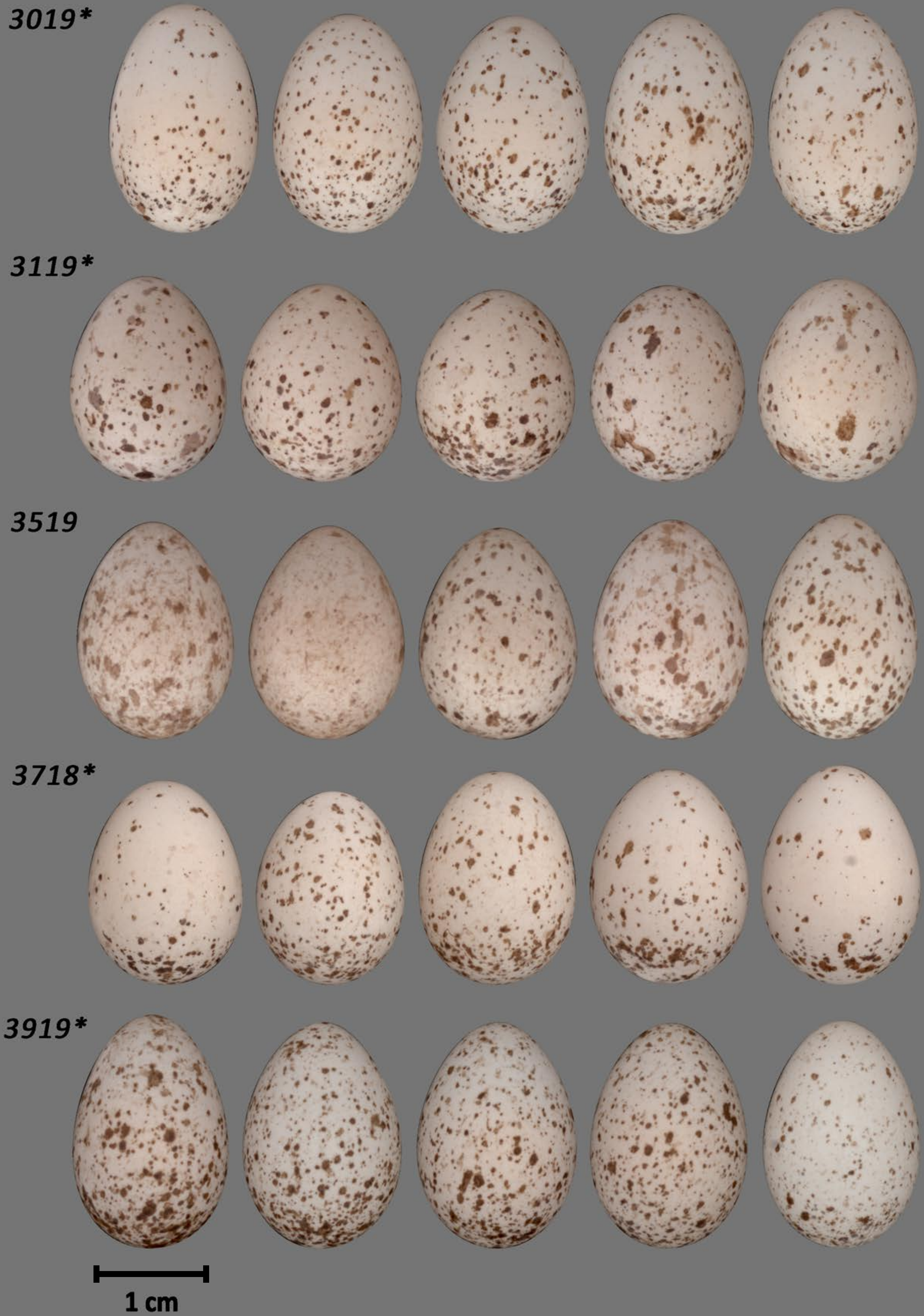
3019*
3119*
3519
3718*
3919*
1 cm

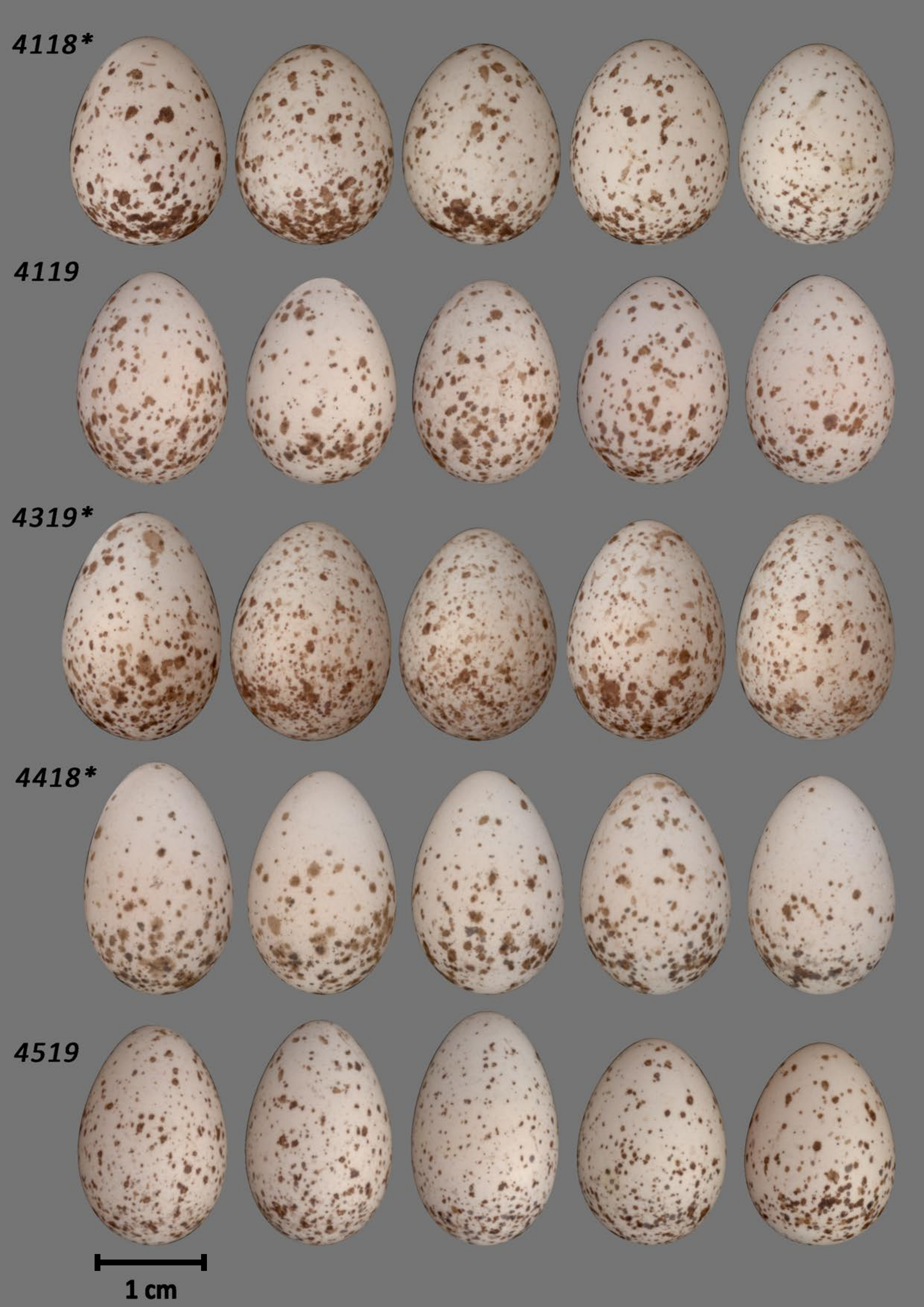
4118*
4119
4319*
4418*
4519
1 cm

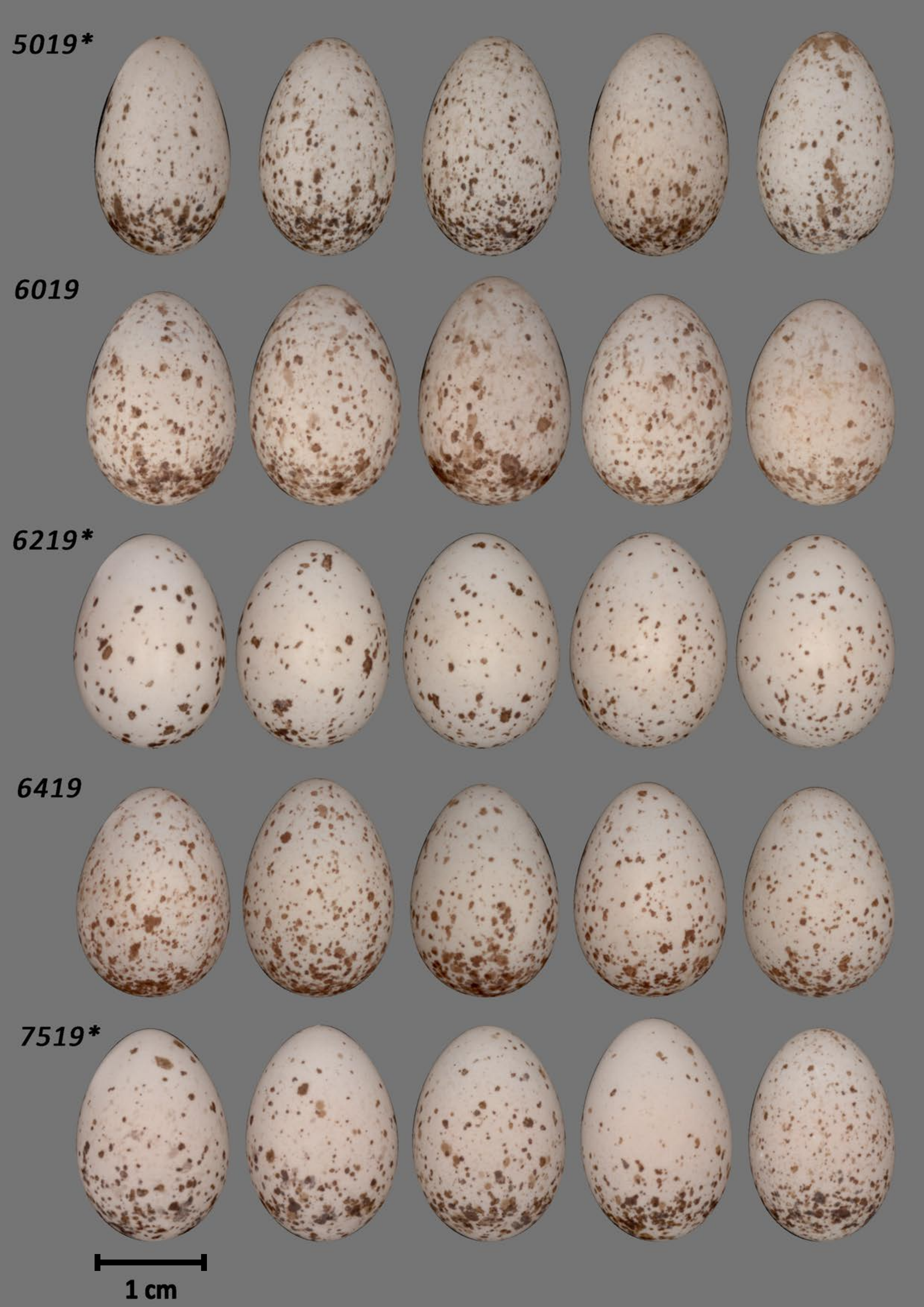
5019*
6019
6219*
6419
7519*
1 cm

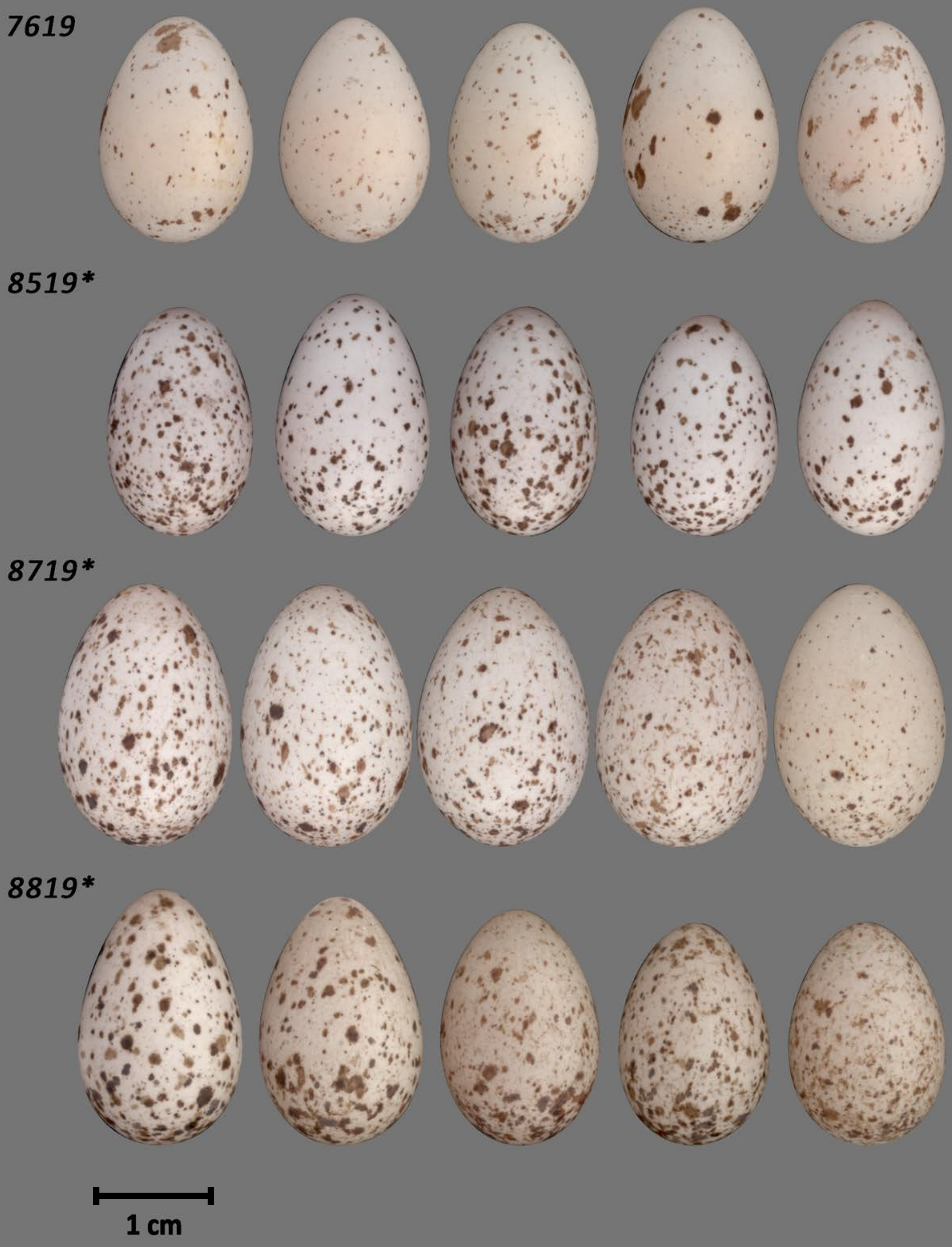
7619
8519*
8719*
8819*
1 cm
